# Multimodal computational framework identifies B cell convergence in autoimmunity and ageing

**DOI:** 10.64898/2026.06.29.735110

**Authors:** Hantao Lou, Meihan Zhang, Bo Zhang, Qianjin Lu, Jianqing Zheng, Xuetao Cao

## Abstract

Identification of the origin of pathogenic immune cells is crucial for therapeutic interventions and diagnosis but pseudotime methods struggle to trace immune cells accurately. Current trajectory inference methods for B cell development and response in health and disease either ignore or underutilize antigen receptor sequence information, limiting their ability to resolve developmental pathways, particularly for pathogenic populations. Widely used methods such as Monocle 3 reconstruct developmental paths from transcriptomic similarity alone, discarding the features from immune receptors. Dandelion has combined the immune receptor features with transcriptomics but it struggles to simulate the trajectory path of B cells. Here we present ClonoTrace, a computational framework that integrates BCR sequence features with transcriptomic trajectory inference through gated fusion of multimodal embeddings. In fetal B cell development and germinal centre development, ClonoTrace achieves higher trajectory inference accuracy than Monocle 3 and Dandelion. Applied to systemic lupus erythematosus, ClonoTrace identifies memory B cell extrafollicular maturation pathway in addition to naïve B cell, accompanied by induction of *ZEB2* with a concomitant decline of *BACH2* along the trajectory, as the alternative origin of pathogenic double negative 2 B cells (DN2) in systemic lupus erythematosus (SLE) patients. In healthy ageing, ClonoTrace identified three pathways from naive, IgM^+^ memory B cells and switched-memory B cells mature through a DN2-associated transcriptional state that precedes age-associated B cells. ClonoTrace’s fate probability algorithm indicated that IgM^+^memory B cell to ABC transition emerged as the leading candidate age-associated transition, that is a process distinct from SLE DN2 maturation. ClonoTrace provides a generalizable framework for receptor-informed trajectory inference, revealing the developmental pathways of pathogenic B cell populations that are untraceable to single modality approaches in autoimmunity and aging.

## 1. Introduction

Understanding how B cells develop, diversify, and commit to effector fates is fundamental to vaccine design, immunotherapy, and autoimmune treatment. Trajectory inference algorithms such as Monocle 3 and CellRank^1,2^ reconstruct developmental paths from single-cell RNA sequencing (scRNA-seq) data, yet purely transcriptomic approaches disregard a defining feature of B cells: their somatically rearranged antigen receptor sequences. SHM burden, class-switch recombination (CSR) state, and clonal expansion provide independent temporal information that could constrain developmental ordering, but existing methods either discard this information or use it only as a clonal label^3^, limiting their ability to accurately capture trajectory state transitions in B cells. B cell development spans a continuum from naïve precursors through germinal centre (GC) reactions to long-lived memory and effector fates^4,5^. Several non-canonical B cell populations have emerged as central players in autoimmunity and ageing. Double-negative 2 (DN2) cells and the related atypical memory (AtM) cells accumulate in systemic lupus erythematosus (SLE) and other autoimmune conditions^6,7^, while age-associated B cells (ABCs), which share *TBX21* and CD11c expression with DN2 cells, expand during ageing and contribute to inflammaging^8–10^. Despite their clinical importance, the pathway hierarchy connecting these populations to their precursors remains unresolved, and their origin is debated. One model posits maturation from naïve B cells through extrafollicular (EF) activation, whereas another supports memory B cell re-differentiation^11,12^. Resolving this debate requires methods that can simultaneously track BCR sequence evolution and transcriptomic fate transitions.

Here we present ClonoTrace, a computational framework that integrates BCR sequence maturation with transcriptomic trajectory inference through gated fusion of multimodal embeddings. By coupling SHM burden, isotype status, and clonal expansion with gene expression similarity, ClonoTrace resolves ambiguities that confound transcriptome-only methods, particularly in settings where cells at distinct maturation stages share similar expression profiles. We benchmark ClonoTrace against established methods across fetal B cell development and GC reactions, then apply it to SLE and ageing cohorts to resolve the memory-versus-naïve origin debate for DN2 cells, define a DN2-to-ABC maturation axis, and characterise age-dependent rewiring of B cell fate allocation.

## 2. Materials and Methods

### 2.1 Datasets

Fetal B cell data comprising paired scRNA-seq and scV(D)J-seq were obtained from published cohorts at https://developmental.cellatlas.io/fetal-immune^13^ yielding 29,818 cells spanning nine developmental stages. Adult lymph node germinal centre data (133,244 cells) from mRNA-vaccinated donors were obtained from vaccinated cohorts^14^. For the SLE cohort, PBMCs were collected from 9 patients with active SLE meeting ACR/EULAR 2019 classification criteria at Xiangya Hospital under institutional ethics board approval; informed consent was obtained from all participants. The healthy ageing cohort comprised 27,801 B cells from 61 donors aged 0–90 years spanning seven B cell populations.

### 2.2 Single-cell library preparation and sequencing

For the SLE PBMC dataset, PBMCs were isolated by density-gradient centrifugation (Ficoll-Paque) within 2 hours of blood draw. Single-cell suspensions were loaded onto the 10x Genomics Chromium Controller targeting 6,000–10,000 cells per sample using the Chromium Next GEM Single Cell 5′ Reagent Kit v2. V(D)J enrichment for BCR sequencing was performed using the Human B Cell VDJ Enrichment Kit. Gene expression libraries were sequenced at ∼30,000 reads per cell on an Illumina NovaSeq 6000 (150 bp paired-end); V(D)J libraries at ∼5,000 reads per cell. Raw FASTQ files were processed using Cell Ranger v7.1.0 with the GRCh38 reference genome (2020-A).

### 2.3 Quality control, normalisation, and integration

Per-cell quality metrics were computed in Scanpy (v1.9). Cells were retained if total counts fell between 200 and the 99th percentile, detected genes between 200 and the 99th percentile, and mitochondrial fraction <20%. Doublets were removed using Scrublet (v0.2.3; score threshold 0.25). Count matrices were normalized to 10,000 counts per cell and log1p-transformed. Highly variable genes (HVGs; n = 3,000) were identified using the Seurat v3 method, supplemented by forced inclusion of 36 curated B cell markers. Multi-sample integration was performed using scVI (v0.20; latent dimension 30, negative binomial likelihood, up to 400 epochs with early stopping). UMAP embeddings were computed with umap-learn (v0.5; min_dist = 0.3). Leiden clustering (resolution 0.5–1.2) was applied to the scVI-derived k-nearest neighbour (kNN) graph.

### 2.4 BCR repertoire preprocessing

Cell Ranger V(D)J output was imported using Dandelion (v0.3; preprocessing only). Per-cell BCR metadata extracted for ClonoTrace included: (i) isotype (C-gene call from the dominant productive heavy-chain contig); (ii) SHM rate (V-gene nucleotide mutations normalized by V-gene length); (iii) clone identity (defined by identical V-gene, J-gene, and CDR3 amino acid sequence); and (iv) full-length heavy- and light-chain amino acid sequences. Cells lacking a productive heavy-chain contig were retained in the transcriptomic analysis but received missing values for all BCR-derived features.

### 2.5 ClonoTrace computational framework

#### 2.5.1 BCR sequence embedding

Full-length heavy- and light-chain amino acid sequences were embedded using AntiBERTy^15^, a transformer language model pre-trained on 558 million unpaired antibody sequences. Per-sequence embeddings were obtained by mean-pooling across token hidden states (H = 512). Heavy and light chain embeddings were concatenated and dimensionality-reduced by PCA to 256 components, yielding X_BCR_ ∈ ℝ^n×256^. Cells lacking light-chain contigs received zero-padded light-chain embeddings.

#### 2.5.2 BCR maturation quality score (q-score)

ClonoTrace assigns each cell a continuous BCR maturation quality score *q_score_* ∈ [0, 1] by integrating a technical BCR confidence score *q_tech_* and a biologically informed maturation score *q_bio_* . Stage 1 computes *q_tech_* from six contig-level sub-metrics (assembly confidence, barcode accuracy, productive rearrangement, full-length coverage, CDR/FWR completeness, and sequencing depth). Stage 2 computes *q_bio_* from three evidence streams: class-switch recombination (isotype), somatic hypermutation (SHM), and clonal expansion. It uses a weighted mean over available components. By default, we use weights *w_iso_*= 0.45, *w_shm_*= 0.45, and *w_clonal_*= 0.10 to balance discrete isotype information with continuous SHM and clonality signals. In some analyses reported in this study, we instead used *w_iso_*= 0.05, *w_shm_*= 0.90, and *w_clonal_*= 0.05, because SHM provided the most informative and robust continuous proxy of antigen experience in these datasets, whereas isotype and clonal size were either missing for a subset of cells or less discriminative across the relevant developmental states. These weights are user-adjustable and should be tuned to the data characteristics (e.g., isotype resolution, clone calling quality, and cohort composition). The resulting *q_bio_* was gated by *q_tech_*. Stage 3 fuses *q_tech_* and *q_bio_* using data-adaptive weights. We first optionally applied a monotonic contrast transform Φ(·) to *q_bio_* , by default a square-root stretch which amplifies separation in the low-scoring range without altering rank order.

The default weights *w_iso_*= 0.45, *w_shm_*= 0.45, and *w_clonal_*= 0.10 were set based on established B cell biology. SHM and class switching represent the gold-standard BCR maturation readouts: both are irreversible, progressive events accumulating monotonically during antigen-driven responses.

SHM reflects cumulative activation-induced deaminase activity; class switching is directional (IgM/IgD to IgG or IgA) and permanent. These features were therefore weighted equally and dominantly (0.45 each). In contrast, clonal expansion is a weaker temporal signal. Expansion can occur at any developmental stage, including early transitional cells, memory recall, and terminal plasma bursts. Clonal expansion is sensitive to sampling depth and batch effects, which was therefore down-weighted (0.10) to limit technical noise while retaining its contribution. Sensitivity analyses addressed two complementary concerns. First, we tested the choice of q-score component weights across a 7-setting ablation set covering the four simplex corners (baseline, iso-dominant, SHM-dominant) plus three component-isolation modes (q_iso_only, q_shm_only, q_clonal_only) and a q-score-neutralised control (no_q) that included the reviewer-flagged SHM-dominant corner (W_shm = 0.60) and iso-dominant corner (W_iso = 0.60) in addition to the baseline (0.45, 0.45, 0.10) and four symmetric perturbations (Supplementary Table 1); reference-cell-type orderings were also varied across four plausible alternatives per dataset (e.g. DZ to LZ swap in the GC benchmark). Composite scores varied across q-score weights by less than the inter-method gap and were essentially invariant to reference-ordering choice.

The final *q_score_* combines *q_tech_* and transformed *q_bio_* using data-adaptive weights *α* and *β* reflecting their effective information content:

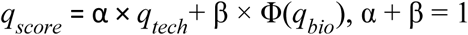

where *α* and *β* is determined by comparing distribution informativeness. For each component, effective information is quantified as:

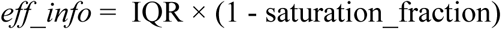

where IQR is inter-quartile range and saturation_fraction is the proportion of cells exceeding a predefined saturation threshold.. Higher IQR and lower saturation indicate greater biological variation. The adaptive weights are defined as:

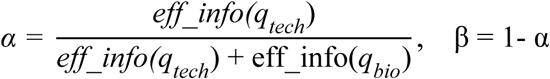

When one component dominates (e.g., *α* > 0.9 or *β* > 0.9), *q_score_* is set to that component alone. This ensures the component with more variation receives proportionally higher weight. For instance, when BCR contigs are uniformly high-quality ( *q_tech_* saturated), *q_bio_* automatically dominates (α → 0).

The weighting scheme, contrast-stretching function, and saturation threshold can be adjusted as needed for different datasets and analytical settings.

#### 2.5.3 Lineage skeleton construction and pseudotime

A cluster-level connectivity graph was constructed from the scVI kNN graph, with edge weights adjusted by median q-score between cluster pairs. Forward edges (increasing q-score) were boosted by (1 + Δq) and backward edges penalised by (1 − |Δq|) before minimum spanning tree (MST) extraction using Kruskal’s algorithm. The MST serves as a lineage skeleton encoding coarse cluster-level developmental ordering. Two pseudotime modes are available: geometric pseudotime (tree distances propagated via linear interpolation) and q-score-aware pseudotime (isotonic regression against q-score to eliminate biologically implausible reversals). Geometric pseudotime projects each cell onto the nearest MST edge in scVI latent space and computes its tree distance from the root; this mode preserves the BCR-informed skeleton topology without further q-score adjustment at the cell level. Q-score-aware pseudotime additionally applies isotonic regression of pseudotime against q-score (convex blend, α = 0.3–0.7) to enforce monotonic alignment with BCR maturation.

#### 2.5.4 Transition kernel construction and macrostate analysis

Trajectory inference is performed via a three-kernel architecture. The lineage-prior pseudotime kernel (LPPK) biases transitions towards pseudotime-advancing neighbours with MST-skeleton gating (penalty strength λ_skel_ = 0.5). The BCR-aware kernel (BCRK) constructs an independent kNN graph in the AntiBERTy embedding space and re-weights transitions by logistic q-score gating (β = 10). The connectivity kernel (CONK) is the standard scVI kNN adjacency matrix acting as a regularizer. The combined transition matrix is T_combined_ = 0.50·T_LPPK_ + 0.30·T_BCRK_ + 0.20·T_CONK_. Terminal macrostates and fate probabilities were identified using the Generalized Perron Cluster Cluster Analysis (GPCCA) module in CellRank^2^.

### 2.6 Benchmarking framework

ClonoTrace was benchmarked against Monocle 3, Dandelion, Palantir^16^, DPT^17^, and VIA^18^ using an eight-metric composite framework. The eight metrics and their weights (summing to 100%) are: Kendall τ marker concordance (25%), Spearman ρ cell-type ordering (15%), Branch AUC (15%), Pairwise Cohen’s d separation (15%), Variance explained η² (10%), Pseudotime statistics (10%), Geodesic correlation (5%), and F1-Branches (5%). Per-metric scores were linearly scaled to [0, 100] and combined into a single composite score; when a metric was not computable on a given dataset, weights were renormalised across the remaining metrics. See evaluate_trajectory.compute_composite_score for the canonical implementation. Palantir, DPT, and VIA, which produce pseudotime-only outputs, were evaluated on Spearman ρ and PT Order only. Bootstrap stability of q-score was assessed across 50 resampling replicates (80% subsampling).

### 2.7 SLE-specific and ageing analyses

Extrafollicular (EF) score and GC score were computed as the mean z-scored expression of curated gene sets (EF: *TBX21*, *ITGAX*, *ZEB2*, *FCRL5*, *FGR*, *HCK*; GC: *BCL6*, *AICDA*, *MKI67*, *CXCR4*, *RGS13*). Bridge cells were defined as memory cells with fate probability towards the DN2/AtM terminal state exceeding 0.4. For per-source-celltype transition analyses (e.g. Memory to DN2 vs Naïve to DN2 per-donor enrichment) we computed the single-step transition mass directly from the combined-kernel row-stochastic matrix T_combined, defined as mass(s→t) = mean_{i ∈ src(s)} Σ_{j ∈ tgt(t)} T_combined[i, j]. This estimand is independent of the GPCCA macrostate selection and was adopted because the GPCCA fit identified Memory (B.03.TXNIP+Bm) as a terminal macrostate, which makes fate-probability(Memory to other) identically zero by construction. Transition mass is therefore the appropriate estimand for testing whether Memory cells preferentially flow toward DN2/AtM relative to Naïve cells. The q-score ablation modes used in sensitivity analyses are defined as follows: "no_q" sets q_score = 1.0 for every cell so the BCRKernel q-direction filter is vacuous (note the upstream pseudotime remains derived from scvi features, so no_q controls the q-score but does not eliminate transcriptional maturation information); "q_iso_only" recomputes q_bio with W_iso = 1.0 and W_shm = W_clonal = 0. This removes SHM and clonal expansion from the q-score while retaining isotype. Pairwise Jaccard clonal overlap was computed between all B cell populations. For ageing analyses, per-donor mean transition probabilities were correlated with donor age using Spearman rank correlation; significance of age-dependent kinetic rewiring was assessed by Fisher z-transformation of Spearman ρ values between young (≤30 years) and old (>60 years) donors. All reported p-values are Benjamini–Hochberg corrected unless otherwise stated.

## 3. Results

### 3.1 ClonoTrace is a multimodal framework for B cell trajectory inference

ClonoTrace is a multimodal computational framework that jointly leverages single-cell V(D)J sequencing, single-cell RNA-seq (scRNA-seq), and a BCR maturation quality metric to infer B cell developmental trajectories at single-cell resolution (**Fig. 1A**). ClonoTrace addresses a key limitation in existing tools: it treats BCR sequence data not merely as a clonal lineage label but as a continuous, cell-level maturation prior that actively guides trajectory inference, enabling the reconstruction of differentiation routes and terminal cell fates. This may facilitate the identification of therapeutic interventions for B cell-mediated diseases.

**Figure 1.**
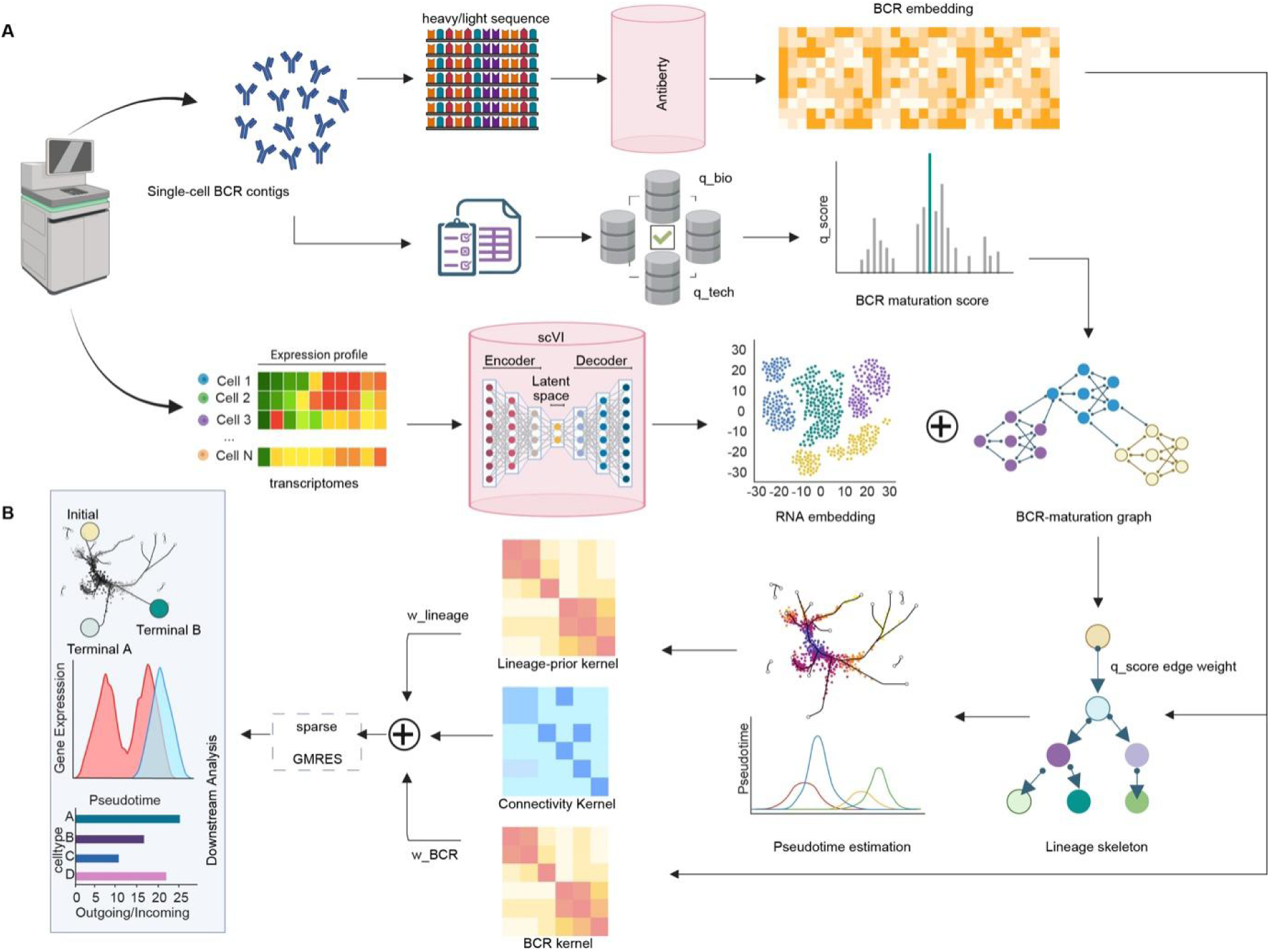
ClonoTrace: a multimodal framework for single-cell trajectory inference for B cell development. (A) Overview of the ClonoTrace pipeline. Paired scRNA-seq and scV(D)J-seq data serve as dual inputs. The RNA modality undergoes standard preprocessing to produce a cell-level kNN graph and UMAP embedding. The BCR modality is processed through two parallel streams: (i) heavy-and light-chain amino acid sequences are embedded via AntiBERTy into a 256-dimensional representation after mean-pooling and PCA reduction; (ii) BCR metadata (isotype, SHM level, clonal expansion) are combined into a composite quality score (q-score). Q-score computation. Three evidence streams weighted by biological informativeness: isotype class-switch status (*w_iso_*, = 0.45), SHM level (*w_shm_* = 0.45), and clonal expansion (*w_clonal_* = 0.10). These are combined into a biological maturity score (*q_bio_*) gated by a technical completeness metric (*q_tech_*). The final q-score is combined of *q_tech_* and *q_bio_* . AntiBERTy BCR sequence embedding architecture. (B) BCR-maturation-reweighted cluster graph and minimum spanning tree (MST) extraction, forming the lineage skeleton for pseudotime computation. Triple CellRank kernel architecture: lineage-prior pseudotime kernel (LPPK) , BCR-aware kernel (BCRK) and connectivity kernel (CONK) are linearly combined into T_combined. Representative application contexts: fetal bone marrow, adult lymph node, SLE PBMCs, and healthy ageing cohort.

To obtain a continuous sequence-level representation of BCR diversity, ClonoTrace embeds full-length BCR heavy and light chain amino acid sequences using AntiBERTy (**Fig. 1A**), a transformer language model pre-trained on 558 million unpaired antibody sequences^13^.

AntiBERTy encodes each chain into a contextual embedding that captures framework and complementarity-determining region (CDR) structural features; these are supplemented by CDR3 physicochemical descriptors (see Methods). To complement language-model representations, ClonoTrace additionally computes CDR3 physicochemical features including hydrophobicity, net charge, aromaticity, amino acid composition, and k-mer frequencies.

Together, the combined BCR embedding captures sequence-level variation in SHM load, isotype usage, and CDR3 biochemistry, providing biologically interpretable features for trajectory inference and differential analysis.

In parallel, ClonoTrace applies scVI^14^ to the scRNA-seq count matrix, learning a low-dimensional latent representation that captures major B cell states while accounting for technical noise and batch variation (**Fig. 1A**). Cell populations are identified by Leiden clustering in this space and annotated by canonical marker gene expression. A cluster-level connectivity graph is then constructed from the scVI latent space using k-nearest neighbor (kNN) graphs, and edge weights are adjusted by the median q-score of connected cells, such that transitions through cells with higher BCR maturation evidence are preferentially retained. Starting from ClonoTrace defined root states by q-score or user-defined early B cell populations as root states, ClonoTrace extracts a minimum spanning tree (MST) from the BCR-maturation-reweighted cluster graph (**Fig. 1A**). This MST functions as a lineage skeleton that encodes an initial cluster-level ordering along putative developmental branches, constraining subsequent cell-level inference to biologically coherent paths. Pseudotime is computed along the skeleton and propagated to individual cells, with optional q-score-guided monotonicity enforcement that reduces pseudotime reversals in cells with low BCR maturation evidence.

Trajectory inference is performed by constructing three complementary CellRank^2^ transition kernels that operate in distinct feature spaces and encode distinct biological principles (**Fig. 1B**). The lineage-prior pseudotime kernel (LPPK) biases cell–cell transitions towards transcriptionally similar neighbors that advance along the MST skeleton in pseudotime, combining a logistic soft-threshold on pseudotime directionality with cluster-level skeleton gating controlled by the penalty strength parameter λ_skel_. Transitions between cells that traverse cluster–cluster edges absent from the MST are penalized, enforcing the coarse lineage topology derived from BCR-maturation-informed graph reweighting. The BCR-aware kernel (BCRK) constructs an independent kNN graph in the AntiBERTy embedding space and re-weights transitions towards cells with higher qscore values using either a hard threshold or a logistic soft-constraint (parameter β), ensuring that cells with immature or low-confidence BCR sequences do not act as spurious attractors in trajectory space. The connectivity kernel (CONK) is defined as the standard scVI kNN adjacency matrix and serves as a regularizer. The three kernels are linearly combined as Kcombined = *α*_1_·KLPPK + *α*_2_·KBCRK + *α_3_*·KCONK, where *α*_1_ + *α*_2_ + *α_3_* = 1 controls the relative contribution of BCR versus transcriptomic evidence. The combined transition matrix is analyzed with the Generalized Perron Cluster Cluster Analysis (GPCCA) module^16^ to identify macrostates, estimate cell fate probabilities towards user-defined terminal populations, and recover driver transcription factors and gene regulatory programs associated with each developmental branch. This modular, kernel-based design is intended to allow ClonoTrace to be applied to diverse BCR-seq and scRNA-seq datasets, as demonstrated below in fetal, lymph node, and peripheral blood contexts (**Fig. 1B**).

### 3.2 ClonoTrace recapitulates canonical fetal B cell maturation and outperforms state-of-the-art methods

To evaluate whether multimodal integration improves concordance with the canonical reference ordering, we benchmarked ClonoTrace against five widely used methods including Monocle 3^1^, Dandelion^3^, Palantir^15^, DPT^17^, and VIA^18^. We gathered a pan-fetal B cell dataset comprising paired scRNA-seq and scV(D)J (**Fig. 2A**). The fetal B developmental hierarchy is well-established, in which Pre-pro-B and Pro-B progenitors through Large pre-B, Small pre-B, Immature B, and Mature B intermediates before committing to either B1 or Plasma B terminal fates^19,20^. Across all eight evaluation dimensions, ClonoTrace achieved the highest performance, with a composite score of 95.2/100 , compared to 76.2/100 for Monocle 3 and 37.3/100 for Dandelion (**Fig. 2B, Supplementary Fig. 1A**). The improvement was most pronounced for global ordering fidelity, which is evaluated by branch AUC and F1 score (**Supplementary Fig. 1A**).

**Figure 2.**
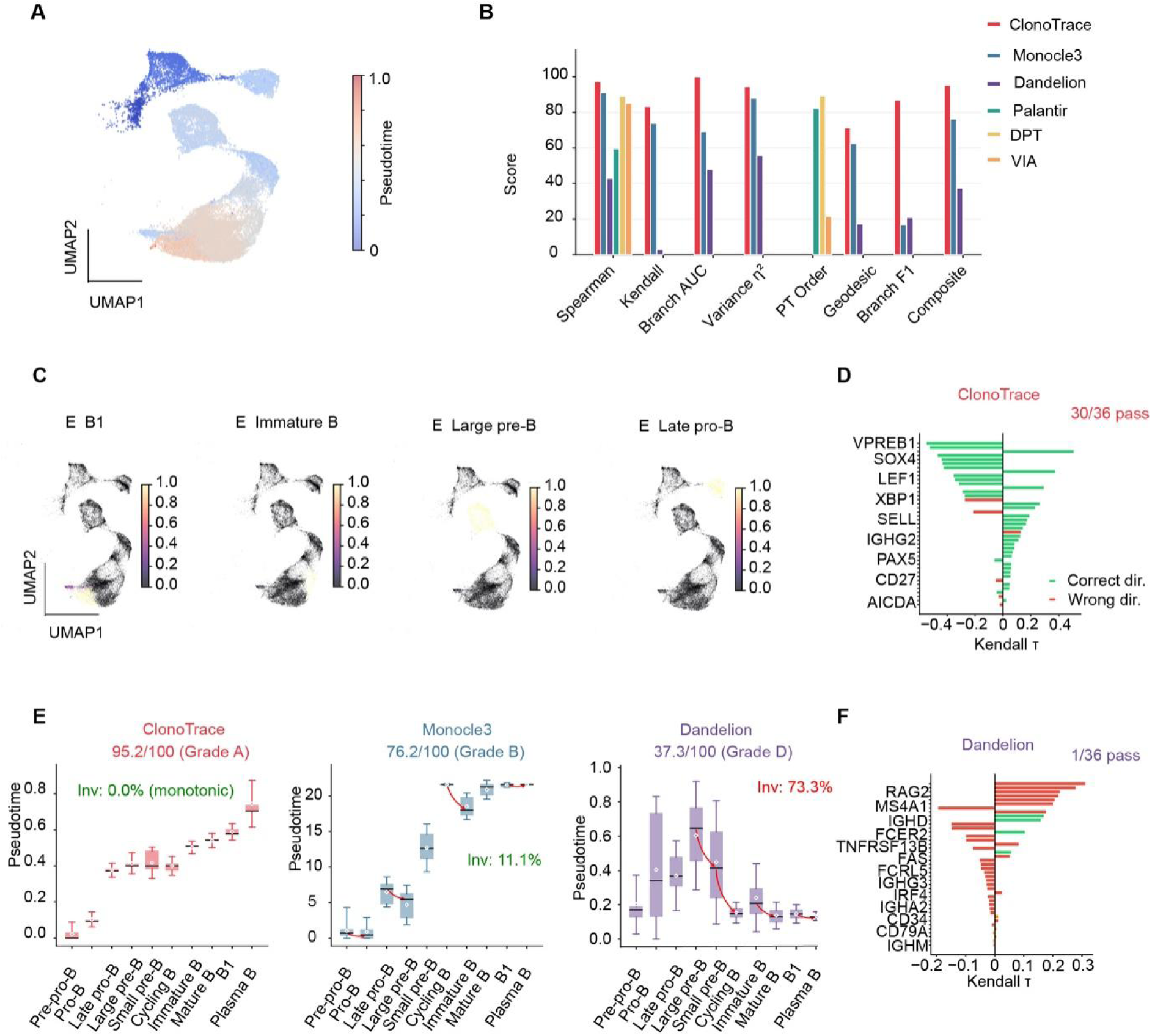
ClonoTrace achieves accurate trajectory inference of fetal B cell development. (A) UMAP embedding of 29,818 fetal B cells coloured by ClonoTrace-inferred pseudotime. The inferred gradient recapitulates the canonical hierarchy: Pre-pro-B → Pro-B → Late pro-B → Cycling B → Large pre-B → Small pre-B → Immature B → Mature B → B1/Plasma B, with 0.0% developmental inversions. (B) Quantitative comparison of six trajectory inference methods across eight evaluation metrics. (C) ClonoTrace-inferred fate probabilities for four representative terminal lineages: B1 (n = 2,897), Immature B (n = 1,106), Large pre-B (n = 4,702), and Late pro-B (n = 2,887). (D) Per-gene Kendall τ between marker gene expression and ClonoTrace pseudotime. Green = correct direction; red = incorrect direction. ClonoTrace passes 30/36 markers (83.3%). (E) Boxplots of pseudotime per cell type in developmental order showing ClonoTrace (0.0% inversions; Spearman ρ = 0.988), Monocle 3 (11.1% inversions), and Dandelion (73.3% inversions). (F) Per-gene Kendall τ for Dandelion.

Visualization of pseudotime on the UMAP embedding confirmed these quantitative findings (**Fig. 2A, Supplementary Fig. 1B**). ClonoTrace produced a monotonically increasing pseudotime gradient (0% inversion rate) from early progenitors in the Pre-pro-B/Pro-B region to terminal Plasma B cells, with each intermediate population occupying a coherent, non-overlapping pseudotime band. In contrast, Monocle 3 generated a less resolved gradient with partial ordering failures near the Immature B to Mature B transition, and DPT showed compressed pseudotime distributions in late-stage populations (**Supplementary Fig. 1B**). Fate probability maps further showed that ClonoTrace resolved four biologically interpretable terminal states that are composed of B1, Immature B, Large pre-B, and Late pro-B with spatially restricted probability distributions consistent with their anatomical and functional positions in the B cell hierarchy (**Fig. 2C**).

To evaluate whether the inferred pseudotime axis captures the correct directionality of molecular programs, we computed Kendall’s τ between pseudotime and expression for 36 canonical fetal B cell marker genes spanning progenitor, intermediate, and mature states (**Fig. 2D, F, Supplementary Fig. 1F**). For each gene, we defined the expected sign of τ based on established expression patterns: progenitor markers (*VPREB1, SOX4, LEF1, RAG2, EBF1, CD79A*) should show negative τ (downregulation along developmental progression), whereas maturation markers (*IGHM, IGHG1/3, CD27, PAX5*) should show positive τ (upregulation with maturation).

ClonoTrace classified concordantly with the expected direction 30 out of 36 markers (**Fig. 2D**), capturing downregulation of progenitor gene expression including *VPREB1* (τ = −0.55), *SOX4* (τ = −0.44), and *LEF1* (τ = −0.35) coupled with progressive upregulation of late B cell and isotype maturation markers. The six discordant genes (*TCL1A, CD27, SDC1, XBP1, MZB1*, and *AICDA*) reflect genes with characteristically non-monotonic expression patterns across the fetal B cell axis **(Fig. 2D, F)**. Dandelion, by contrast, classified concordantly with the expected direction only 1 of 36 markers (**Fig. 2F**), with 11 markers showing directionally incorrect directional τ values.

ClonoTrace achieved the highest Spearman ρ and Kendall τ with highly stable q-score (**Supplementary Fig. 1D, E**), reflecting close alignment with the canonical Pre-pro-B to Pro-B, followed by Large pre-B, Small pre-B, Immature B, Mature B and B1 to Plasma B hierarchy (**Figure. 2E**). The inversion rate, defined as the fraction of adjacent cell-type pairs for which pseudotime order is reversed relative to the canonical reference, was 0.0% for ClonoTrace (fully monotonic), compared to 11.1% for Monocle 3 and 73.3% for Dandelion (**Fig. 2E**). These results demonstrate that ClonoTrace produces a pseudotime ordering that is globally consistent with established B cell developmental biology.

### 3.3 ClonoTrace recapitulates germinal center B cell maturation pathways

To assess whether ClonoTrace could map the more complicated non-linear B cell maturation scenario, we next applied it to adult lymph node (LN) B cells undergoing active germinal center (GC) reactions. We profiled adult LN B cells^30^ by paired scRNA-seq and scV(D)J-seq and resolved 13 transcriptionally distinct populations (**Supplementary Fig. 2A, left**). A Pre-GC cluster expressing activation markers but lacking canonical GC zonation markers served as the putative GC-entry state. The GC itself was resolved into canonical DZ-like (dark zone: CXCR4⁺, MKI67⁺, AICDA⁺) and LZ-like (light zone: CD86⁺, BCL6⁺, FDC-interaction signatures) populations, consistent with the canonical GC reaction cycle. The BCR q-score was distributed in a biologically coherent gradient across this landscape (**Supplementary Fig. 2A, right**), rising from near-zero values in naïve clusters through intermediate levels in Pre-GC and memory cells, to maximal values in LNPC, which is characterised by the highest SHM burden and most extensive class switching in the dataset. This gradient was visually distinct from the pseudotime gradient (**Supplementary Fig. 2A, center**), confirming that q-score provides additional features to the transcriptomic pseudotime ordering.

We benchmarked ClonoTrace against Monocle 3 and Dandelion on the LN GC dataset. The reference ordering used for Spearman and Kendall metrics was defined as TCL1A⁺ Bn / NR4A2⁺ Bn, Pre-GC, DZ-like, LZ-like, EGR1⁺ Bm / GPR183⁺ Bm / TXNIP⁺ Bm, LNPC, reflecting the established sequence of GC entry, affinity maturation, exit, and plasma cell commitment^4,5^.

ClonoTrace achieved a composite score of 79.0/100, outperforming Dandelion at 46.1/100 (**Fig. 3A, B**). The inversion rate for ClonoTrace was 14.1% compared to 39.7% for Dandelion, a 2.8-fold reduction in ordering errors (**Fig. 3A, B**). The GC reaction exhibits complex, non-monotonic gene expression dynamics that challenge simple pseudotime ordering. Genes such as *BCL6* and *AICDA* cycle with dark zone/light zone oscillations, violating the monotonic assumption of linear pseudotime. This explains the reduced marker concordance across all methods and highlights a fundamental limitation of pseudotime-based inference while ClonoTrace is more reliable in capturing the maturation pathways in cyclical biological processes.

**Figure 3.**
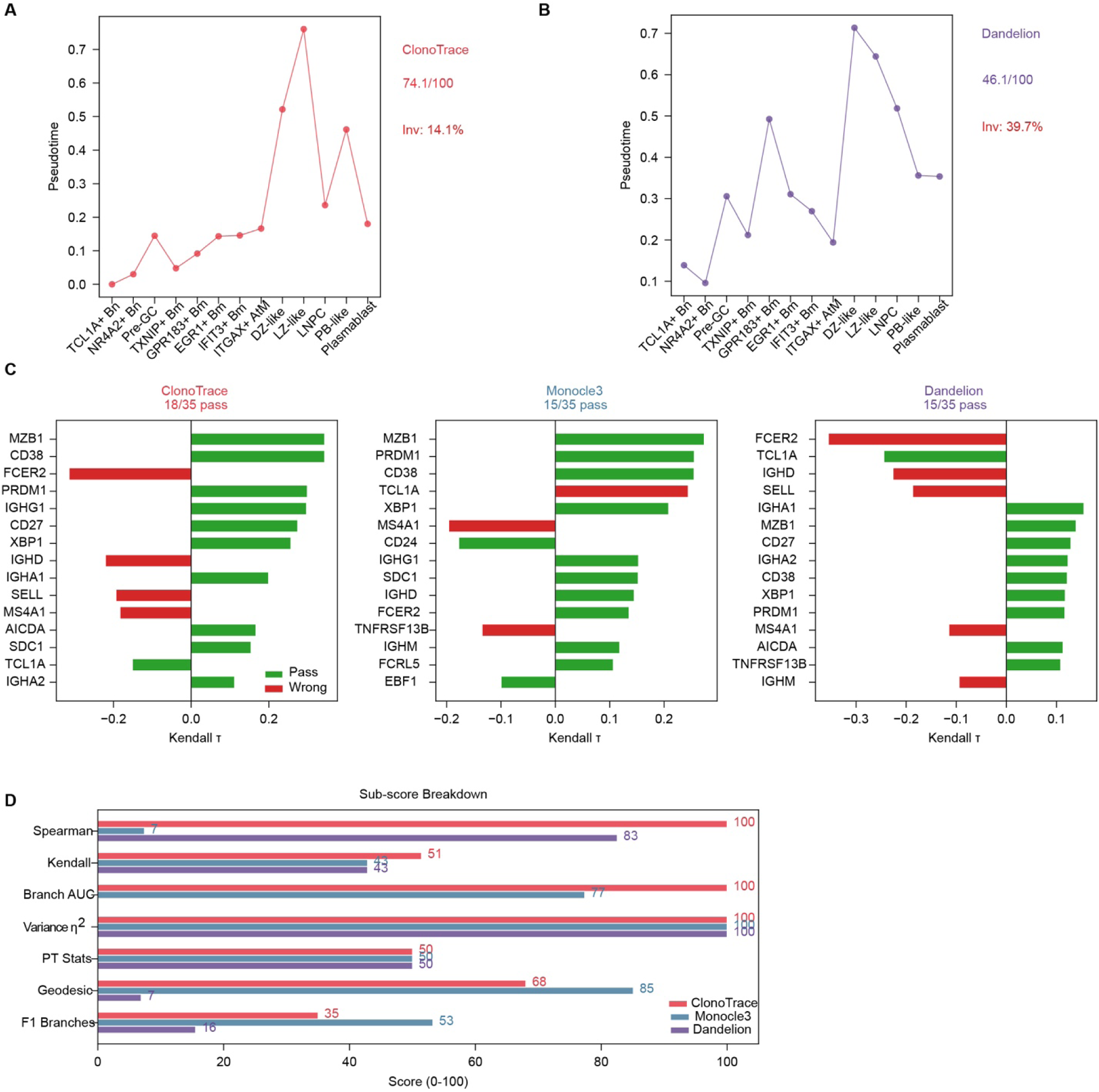
ClonoTrace maintains concordance with the canonical reference ordering and outperforms Dandelion and Monocle 3 on germinal center B cell dataset. (A) ClonoTrace pseudotime ordering of 133,244 GC B cells spanning 13 B cell types. Inversion rate: 14.1%. (B) Dandelion pseudotime on the same GC dataset. Inversion rate: 39.7%. Inversions are coloured in red. (C) Per-gene Kendall τ concordance for ClonoTrace (18/35 pass, 51.4%), Monocle 3 (15/35, 42.9%), and Dandelion (15/35, 42.9%). Green = pass (correct direction, BH p < 0.05); yellow = correct but non-significant; red = wrong direction. (D) Quantitative comparison across trajectory accuracy metrics for ClonoTrace, Monocle 3, and Dandelion. Separation metric scored 0 for ClonoTrace and Dandelion due to mismatched expected pairs (fetal B framework applied to GC context), not method failure.

Marker gene directional validation by Kendall τ per gene revealed a substantial improvement by integrating BCR embedding into the pseudotime algorithm (**Supplementary Fig. 2B**). ClonoTrace classified concordantly with the expected direction 18 of 35 canonical LN B cell markers (51.4%), ahead of both Monocle 3 and Dandelion (each 15/35, 42.9%). Genes that do provide clear directional signals in the GC reaction, including *ITGAX* and *LEF1* as maturation markers, and *DNTT* and *RAG2* as progenitor markers were correctly captured by ClonoTrace (**Fig. 3C, left panel**). These results indicated that ClonoTrace correctly captures the directional structure of the trajectory for genes with monotonic expression patterns along the GC maturation axis.

Examining the eight evaluation metrics individually revealed that ClonoTrace and Monocle 3 exhibited complementary strengths and weaknesses in the LN context (**Fig. 3D**). ClonoTrace achieved the highest Spearman ρ (100 versus 83 for Monocle3 versus 7 for Dandelion), indicating that the cluster-level median pseudotime ordering was near-perfectly aligned with the canonical reference (**Fig. 3D**). ClonoTrace also attained perfect Branch AUC (100) compared to 77 for Monocle 3, a metric that specifically measures how accurately fate probabilities assign cells to their correct developmental branches (**Fig. 3D, Supplementary Fig. 2C**). Bootstrap resampling validated the reproducibility of q-score computation in the LN GC context: across 50 resampling replicates, the Spearman ρ between q-score and expression rank showed a mean of 0.9328 with a standard deviation of 0.0003 and a 95% confidence interval of [0.9322, 0.9334] (**Supplementary Fig. 2D**). Sensitivity analysis of trajectory performance to the choice of q-score contrast-stretching method confirmed that root_g2 is the optimal default for LN GC datasets: Kendall τ = 0.493, the highest value across all methods tested (**Supplementary Fig. 2E**). These results demonstrated that the terminal state identification benefits from being constrained by BCR maturation across datasets (**Supplementary Fig. 2 B, F**).

### 3.4 Clonotrace identified a memory B cell to double negative B cell pathway

The DN B cells have been found to be one of the key pathogenic autoreactive B cell subsets in systemic lupus erythematosus, while the origin and developmental pathway of this population remains elusive. We tackled this challenge with ClonoTrace. We profiled peripheral blood mononuclear cells (PBMCs) from 9 patients with SLE by paired scRNA-seq and scV(D)J, resulting in five main B cell subsets, including naïve B cells, IFN-induced B cells, class-switched memory B cells, double negative B cells, and plasmablasts (**Fig. 4A, Supplementary Fig. 3A**). Two naïve-like clusters were presented as naïve B cells and IFN-naïve B cells, the latter characterised by elevated interferon-stimulated gene expression consistent with the type I interferon signature that distinguishes a subset of circulating naïve B cells in SLE^6,7^(**Supplementary Fig 3A, B**). Three more mature clusters were evident, consisting of a memory compartment , a DN2 memory cluster, co-expressing *TBX21*, *FCRL5*, *LILRB2*, and *SIGLEC6* (**Supplementary Fig. 3A**).

**Figure 4.**
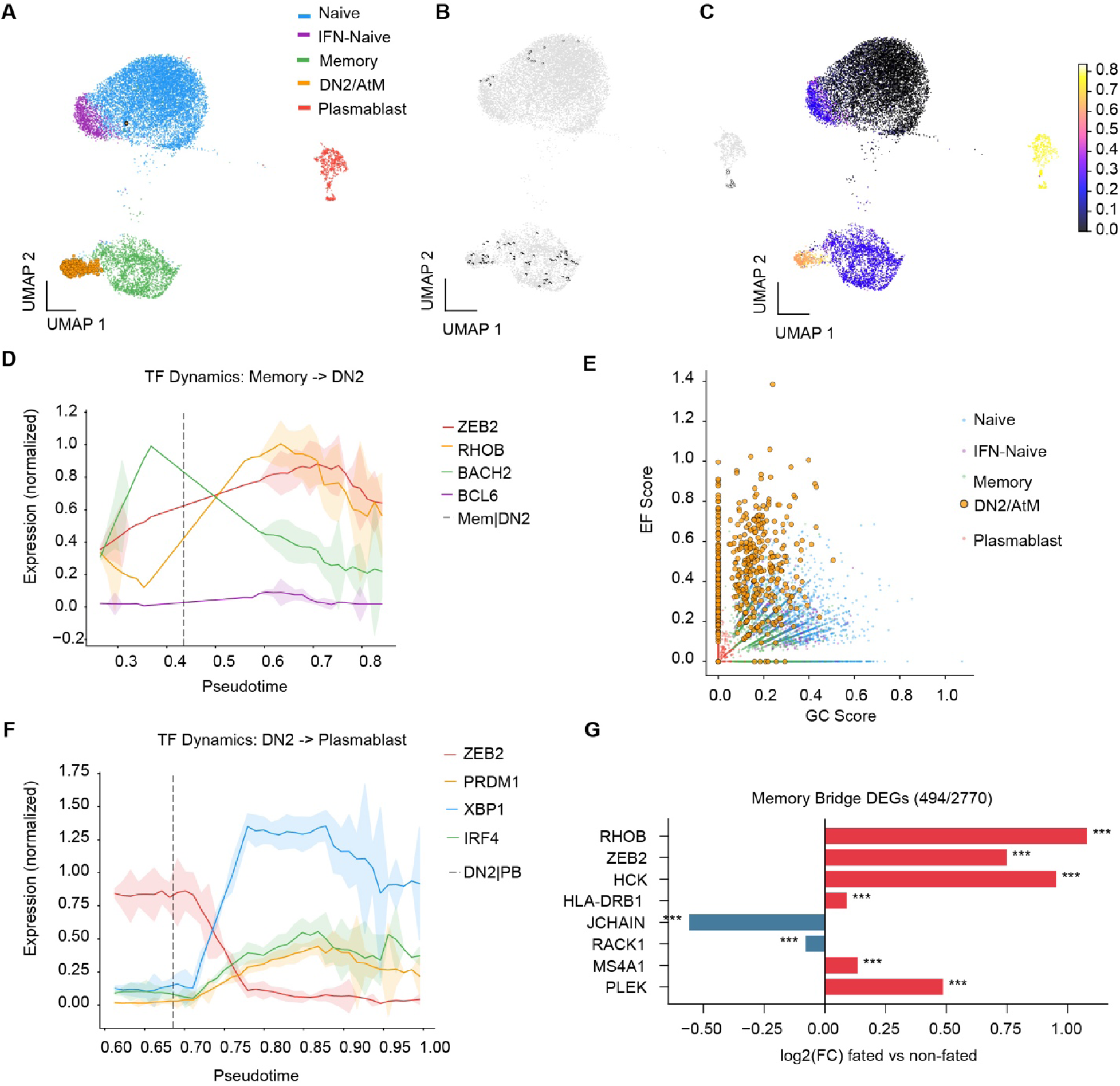
ClonoTrace resolves DN2/atypical memory B cell differentiation trajectories in SLE patients. (A) UMAP embedding of 14,637 B cells from SLE patients (n = 9) coloured by five cell types: Naïve (TCL1A⁺ Bn), IFN-Naïve (IFIT3⁺ Bn), Memory (TXNIP⁺ Bm), DN2/AtM (ITGAX⁺ atypical memory), and Plasmablast. DN2/AtM cells (n = 473, 3.2%) are highlighted. (B) GPCCA-identified macrostate terminals on the UMAP. (C) UMAP coloured by ClonoTrace-inferred pseudotime (viridis scale, 0–0.8). Mean pseudotime per cell type: naïve B, 0.010; IFN-naïve B, 0.275; memory B, 0.259; DN2/AtM, 0.646; plasmablast, 0.793. (D) Transcription factor dynamics along the Memory to DN2 transition: *ZEB2* (red), *RHOB* (yellow), *BACH2* (green), and *BCL6* (purple) expression along pseudotime. The grey dashed line marks the Memory-to-DN2 transition point. Shaded regions indicate SEM. (E) EF score versus GC score per cell, coloured by cell type. DN2/AtM cells cluster exclusively in the EF⁺GC⁻ quadrant. (F) Transcription factor dynamics along the DN2 to Plasmablast transition: *ZEB2* (purple), *XBP1* (blue), *PRDM1* (yellow), and *IRF4* (green). (G) Box plot comparing SHM levels between DN2-fated memory cells (n = 494) and non-fated memory cells (n = 2,276). DN2-fated memory cells display significantly lower SHM (Cohen’s d = −0.23, Mann–Whitney p < 0.001).

To identify terminal B cell fates in an unsupervised manner, we applied the CellRank GPCCA module to the BCR-aware Markov transition matrix (**Fig. 4B**). GPCCA resolved a small number of terminal states, each with stability index SI > 0.96, spanning across the major B cell subsets (**Figure 4B**). Mapping per-cell fate probabilities into a four-vertex fate simplex provided a continuous, multi-resolution view of fate commitment across the B cell landscape (**Fig. 4C**).

Pseudotime imposed a continuous temporal ordering across these populations and memory B cells demonstrated high transition probability to DN2 (**Supplementary Fig. 3C**). A subset of Memory and DN2/AtM cells displayed markedly elevated SHM burden and class-switched isotypes, while the bulk of Naïve cells are significantly lower in SHM, consistent with minimal antigen experience (**Supplementary Fig. 3D**). Further investigation into the transcriptional program that drives this transition, we examined key transcription factor (TF) dynamics along the Memory to DN2/AtM branch (**Fig. 4D**). In the Memory compartment (pseudotime ∼0.25–0.40), *BACH2* was expressed at its highest level and then declined sharply at the branch point (pseudotime ≈ 0.40). Coinciding precisely with this *BACH2* downregulation, *ZEB2* and *RHOB* rose abruptly and were sustained throughout the DN2/AtM compartment (pseudotime 0.40–0.85). *ZEB2* directly promotes *ITGAX* and *FCRL5* expression while suppressing CXCR5-mediated GC entry^22,23^, linking its induction to the acquisition of DN2/AtM surface identity and concurrent GC exclusion. *BCL6* remained near zero throughout the entire Memory to DN2/AtM branch (**Fig. 4D**), providing transcriptional evidence that GC re-entry does not occur. Together, up-regulation of *ZEB2* and *RHOB* in bridge cells (FDR = 2.1×10^−10^ and 5.5×10^−14^, respectively), accompanied by a concomitant decline in *BACH2* along the Memory to DN2 pseudotime trajectory (**Fig. 4D**), mark this transition.We note that *ZEB2* and *RHOB* reach statistical significance as bridge-cell differentially expressed genes, whereas the *BACH2* decline is observed as a pseudotime trend rather than a significant bridge-cell differential expression result. To identify the immediate cellular source of the DN2/AtM fate, we computed mean transition probabilities from each B cell population to the DN2/AtM terminal state using the ClonoTrace Markov transition matrix. Memory B cells showed the highest per-donor transition mass to DN2/AtM, 13.8-fold that of Naïve cells (cell-level pooled ratio 12.30-fold; two-sided paired Wilcoxon p = 0.0039, nine of nine donors; **Supplementary Fig. 3C**; Supplementary Table 2).

These results identify Memory B cells as a major route to DN2/AtM accumulation in this SLE cohort. Scoring each cell for GC activity (*BCL6*, *AICDA*, *MKI67*, *CXCR4*, *RGS13*) and EF activation (*TBX21*, *ITGAX*, *ZEB2*, *FCRL5*, *FGR*, *HCK*) revealed that DN2/AtM cells cluster exclusively in the EF^+^GC^−^ quadrant of the EF–GC biaxial plot (**Fig. 4E**). Memory cells occupied low-to-intermediate EF scores with negligible GC activity, while Naïve and IFN-Naïve cells clustered near the origin (**Fig 4E**).

A key prediction of the EF re-differentiation model is that DN2/AtM-fated Memory cells should carry lower SHM burden than non-fated counterparts. Indeed, DN2-fated Memory cells (fate probability > 0.4) displayed significantly lower SHM rates than non-fated cells (Cohen’s *d* = −0.23, *P* < 0.001, Wilcoxon rank-sum test; **Fig. 4F**), providing an independent BCR-level line of evidence, orthogonal to the transcriptional EF/GC scoring (**Fig. 4E**), that the Memory to DN2/AtM trajectory bypasses GC-mediated affinity maturation. SHM rate along the full pseudotime trajectory showed a continuous increase from the Naïve compartment through Memory to peak values in DN2/AtM and PB cells (**Supplementary Fig. 3D)**. Additionally, the DN2/AtM to PB branch revealed a canonical plasma cell differentiation sequence (**Fig. 4F**). The DN B cell to the antibody-secreting B cells path follows a *ZEB2* decline at the branch point (pseudotime ≈ 0.65), coupled with *PRDM1*, *IRF4*, and *XBP1* upregulation, with *XBP1* reaching peak normalized expression of ∼1.25–1.30, indicating that DN2/AtM cells represent a poised effector state capable of further differentiation into antibody-secreting PBs rather than a transcriptional dead end (**Fig. 4F**). This lower SHM burden in DN2-fated Memory cells reflects truncated or absent GC passage, consistent with extrafollicular activation.

Clonal analysis by Pairwise Jaccard clonal overlap demonstrated that Memory and DN2/AtM cells showed clonal overlap, further supporting the transition between memory B cells to DN2 cells (**Supplementary Fig. 3E, F**). No clonal overlap was detected between DN2/AtM and GC-associated states, providing BCR-lineage-level evidence that the DN2/AtM compartment in SLE is clonally segregated from the GC response. In contrast, only 6 of 1,092 IFN-Naïve cells (0.5%) were identified as DN2/AtM-fated bridge cells, and these showed no significant enrichment for ISG or EF scores compared to non-bridge IFN-Naïve cells (**Supplementary Fig. 3G**), confirming that the IFN-Naïve compartment contributes negligibly to DN2/AtM accumulation despite its interferon activation state. ClonoTrace bridge cell analysis identified 494 of 2,770 Memory cells (17.8%) as committed to the DN2/AtM fate before completing the transcriptional transition. To assess whether this transition is a pooled-cohort artefact or a per-patient signal, we computed combined-kernel transition mass from Memory to DN2/AtM and Naïve to DN2/AtM separately for each of the 9 donors. The per-donor mean Memory to DN2 transition mass (0.0179, averaged over 9 donors) exceeded Naïve to DN2 (0.0013) by a factor of 13.8 (**Supplementary Fig 3I**). Differential expression between bridge and non-bridge Memory cells revealed strong upregulation of *RHOB* (log_2_FC = +1.08), *ZEB2* (+0.75), *HCK* (+0.96), and the MHC-II gene *HLA-DRB1,* mirroring the concomitant transcription factor changes shown (**Fig. 4G**). Conversely, bridge cells downregulated *JCHAIN* which is a hallmark of GC-dependent plasma cell differentiation^24^, provided further molecular evidence against a GC-dependent origin (**Fig. 4G**).

To test whether the Memory-biased entry into the atypical/DN2 state generalises beyond the discovery cohort, we applied ClonoTrace to an independent SLE scRNA-seq + BCR dataset (GSE254176, n = 6 donors), run in an SHM-removed configuration because somatic-hypermutation calls were unavailable (q-weights: isotype 0.8, clonal 0.2). All six donors showed greater Memory to DN2 than Naïve to DN2 transition mass (6/6; one-sided paired Wilcoxon p = 0.016; Supplementary Fig 3J, Supplementary Table 3).

Taken together, the transition probability, EF^+^GC^−^ transcriptional identity and lower SHM burden in DN2-fated Memory cells, and BCR clonal sharing without GC overlap indicate that extrafollicular re-differentiation of memory B cells is the alternative route to DN2/AtM accumulation in SLE.

### 3.5 ClonoTrace delineates three extrafollicular pathways to age-associated B cells

DN2/AtM cells are developmentally linked to age-associated B cells (ABCs), a class-switched, T-bet^+^ FCRL4^+^ population that accumulates progressively with age^8,9,21^. Three routes to ABC differentiation have been proposed based on the transcriptomics and flow cytometric evidence. The three pathways are composed of direct generation from naïve B cells via extrafollicular activation (Naïve to ABC), re-differentiation from switched memory B cells (SwMem to ABC), and from IgM^+^ unswitched memory cells (IgM-Mem to ABC)^10,11^. Whether these pathways are quantitatively equivalent, and how their relative contributions change with age has not been resolved. We applied ClonoTrace to a healthy cohort^19^ (n = 56) spanning donors aged 0–90 years, profiling seven B cell populations: Transitional B, Naïve Resting, IFN-Naïve, IgM Memory, Switched Memory, DN2_preABC, and ABC (**Fig. 5A**).

**Figure 5.**
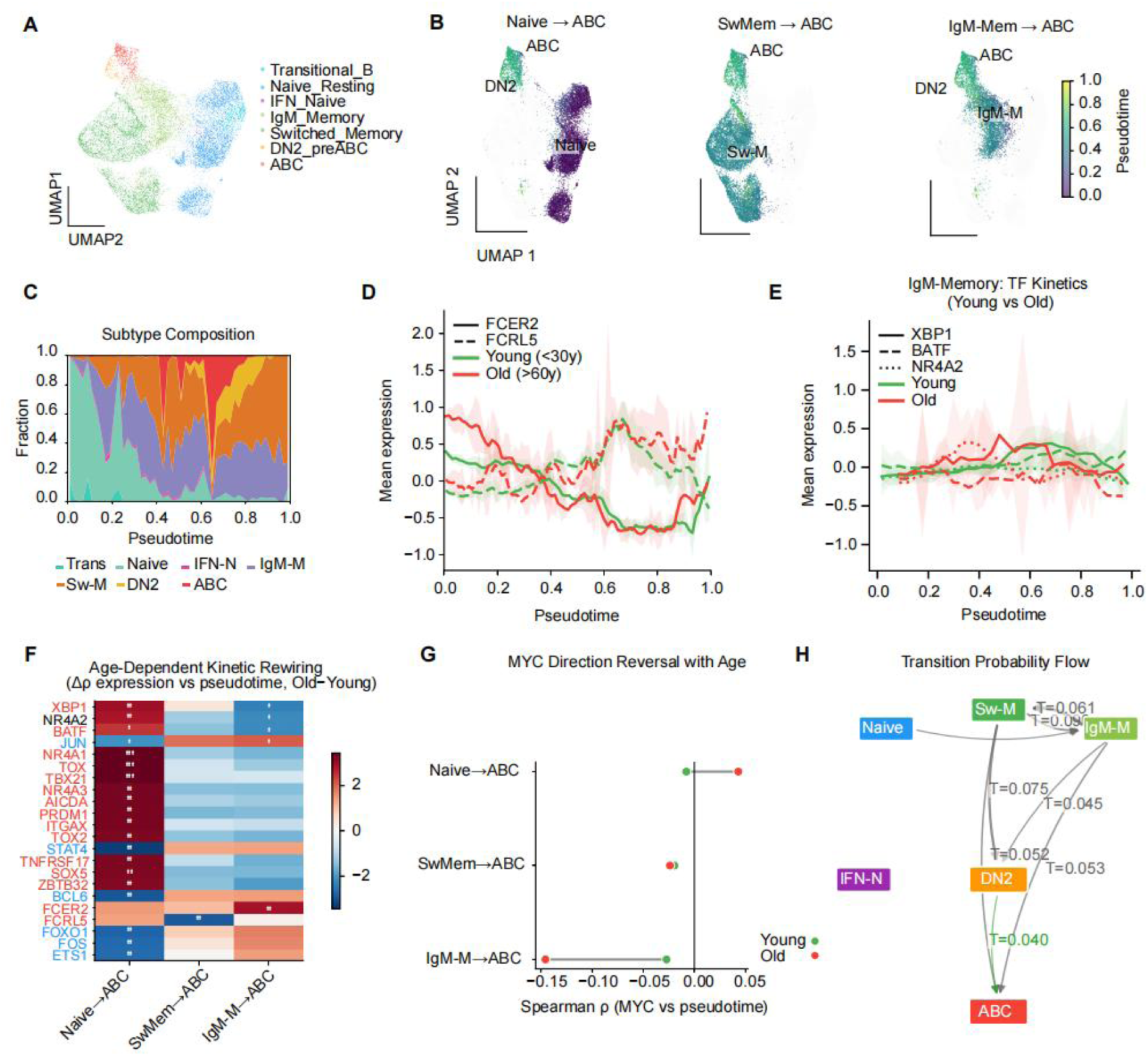
Three transcriptionally similar ABC origin pathways with age-dependent kinetic rewiring. (A) UMAP embedding of 27,801 B cells from 61 donors coloured by seven B cell subtypes: Transitional B, Naïve Resting, IFN-activated Naïve, IgM Memory, Switched Memory, DN2_preABC, and ABC. (B) Three ABC origin pathways visualised on UMAP, coloured by pseudotime gradient. Left: Naïve pathway; Centre: Switched Memory pathway; Right: IgM Memory pathway. (C) Stacked area plot showing subtype composition along pseudotime. Naïve and transitional cells dominate early pseudotime (0.1–0.4); DN2/ABC at intermediate pseudotime and late pseudotime (0.4–0.8); memory B cells emerge at late pseudotime (>0.4). (D) *FCRL5* (dashed) and *FCER2* (solid) expression along pseudotime in young (<30 y, green) and old (>60 y, red) donors. Shaded regions indicate SEM. (E) *XBP1* (solid), NR4A2 (smaller dashed) and *BATF* (dashed) expression along pseudotime in young (<30 y, green) and old (>60 y, red) donors. Shaded regions indicate SEM (F) Age-dependent kinetic rewiring heatmap showing Fisher z-test significance for changes in Spearman ρ between young and old donors across three pathways. (G) Spearman ρ between *MYC* expression and pseudotime in young versus old donors per pathway, showing direction reversal in the IgM-Memory to ABC pathway. (H) Transition probability flow diagram showing DN2 as a convergence point (T = 0.040).

ClonoTrace trajectory inference resolved all three proposed ABC origin pathways as distinct, reproducible pseudotime gradients (**Fig. 5B**). In the Naïve to ABC trajectory, pseudotime increased continuously from Naïve cells through the IFN-Naïve cluster and the DN2_preABC intermediate before terminating in ABCs. The IFN-Naïve compartment occupying a discrete intermediate pseudotime band consistent with an interferon-primed transitional state prior to EF-biased commitment (**Fig. 5B**). In the SwMem to ABC trajectory, switched memory cells (Sw-M) seeded a branch that traversed the DN2_preABC cluster before reaching ABCs, with little contribution from the IFN-Naïve B cells (**Fig. 5B, middle**). The IgM-Mem to ABC trajectory showed the most distinct architecture: IgM-bearing memory cells (IgM-M) are computationally inferred to mature into the DN2 cluster with high pseudotime directionality, bypassing the switched memory states (**Fig. 5B, right**).

Examination of B cell subtype composition as a function of pseudotime confirmed the sequential structure of the trajectory hierarchy (**Fig. 5C**). Naïve cells dominated early pseudotime (0.1–0.3), IFN-Naïve and IgM Memory cells rose at intermediate values (0.3–0.5), DN2_preABC peaked sharply at 0.45–0.9, which overlaps with ABC. Along all three trajectories*, FCER2* and *FCRL5* showed the most significant age-related upregulation along pseudotime (**Fig. 5D**). The induction of *FCER2* was amplified in older donors (>60 years) compared to younger donors (<30 years), initiating at pseudotime 0 to 0.2 for the naïve B cells and IgM^+^ B cells, with peak expression approximately 2.0–2.5-fold higher (**Fig. 5D, Supplementary Fig. 4A**). A broader analysis of nine canonical TFs (*TBX21*, *ZEB2*, *ITGAX*, *TOX*, *TOX2*, *IRF4*, *PRDM1*, *XBP1*, *BACH2*) confirmed the trends. *BCL6* and *IRF4* showed minimal age-dependent ρ changes, confirming that GC-associated and plasma cell maturation remain stable across age groups (**Supplementary Fig. 4B**). In contrary, antibody-secreting B cell genes are downregulated along pseudotime for the IgM^+^ B cells in the older cohorts, XBP1 and BATF expression was found to be significantly lower in the IgM+ B cells at pseudotime higher than 0.7 (**Fig. 5E**). The switched memory B cell to ABC transition revealed a distinct transcriptional entry mechanism by *SOX5* downregulation (log_2_FC ≈ −0.85, p < 0.01) with *NR4A2*, *BATF*, *XBP1*, and *TNFRSF17* upregulation (log_2_FC = +0.8–1.2; **Supplementary Fig. 4C**).

The process of ageing might rewire the kinetics of gene regulation along all three ABC origin trajectories, as revealed by a systematic comparison of gene–pseudotime Spearman correlations between young and old donors across the three pathways (**Fig. 5F, Supplementary Fig. 4D**). In the naïve B cell to ABC pathway, the most dramatic kinetic changes with age included significant increases in *XBP1*(**), *TBX21* (***), *ITGAX* (**), and *NR4A1* (***) correlation with pseudotime(**Fig. 5F, Supplementary Fig. 4E**). Conversely, *XBP1* and *BATF* gene–pseudotime correlations decrease in the IgM^+^ memory B cell to ABC pathway, suggesting that the transcriptional programme coupling IgM^+^ memory B cell maturation into antibody-secreting B cells decreases with age in this pathway (**Fig. 5F**). Together, these results indicate that ageing restructures the regulatory logic of B cell maturation pathways. The unswitched memory route emerged as the most age-related origin of ABC accumulation with weaker antibody secreting ability. We further validated the findings by analysis on a per-donor basis across 52 donors (**Supplementary Fig. 4F)**

Among the age-dependent kinetic changes was a reversal in the direction of MYC–pseudotime correlation specifically in the naïve B cell to ABC pathway (**Fig. 5G**). In young donors, MYC was negatively correlated with pseudotime along this branch (Spearman ρ < 0, * P < 0.05), while the MYC–pseudotime correlation was reversed in the older donors (Spearman ρ > 0, *** P < 0.001), indicating that MYC expression increases progressively with pseudotime along the naïve B cell to ABC branch in aged individuals (**Fig. 5G**).

All three pathways shared transcriptional state with the DN2 like B cell subset before reaching ABCs, and the transition probability from IgM^+^ B cells directly to the ABC increases significantly with age (**Fig. 5H**). Stratifying ABCs by senescence status further defined the functional consequences of DN2-like B cell maturation. The high-senescence ABCs (Hi-sen; n = 709 clones) preferentially originated from DN2-like B cell and Switched Memory compartments, whereas low-senescence ABCs (Lo-sen; n = 708 clones) traced proportionally more to Naïve Resting and unswitched Memory precursors (**Supplementary Fig. 4G**), indicating that transit through the DN2-like B cell is associated with accumulation of senescence-associated transcriptional marks. Hi-sen and Lo-sen ABCs were nearly completely clonally independent (< 1% shared clones; **Supplementary Fig. 4H**), confirming that these represent distinct developmental lineages rather than the same clone at different maturation stages. Transcriptionally, Hi-sen ABCs were selectively enriched for *TBX21*, *ZEB2*, *ITGAX*, and *TOX* relative to Lo-sen ABCs (all p < 0.001), with no significant difference in plasma cell (*PRDM1*, *XBP1*), GC-associated (*BCL6*), or proliferative ( *AICDA*) factors (**Supplementary Fig. 4I**).

### 3.6 Upregulation of unswitched Memory to DN2-like B cell via extrafollicular transition

We further correlated per-donor mean transition probabilities between each pair of B cell populations with donor age across the full cohort, identifying those transitions that are most robustly age-regulated across individuals rather than in aggregate pseudotime comparisons. Two transitions emerged as the strongest age-associated changes with opposite directionality. The transition from unswitched Memory B to DN2-like B cell was the most significantly age-upregulated transition in the dataset (Spearman ρ = 0.551, p = 2.3×10^−5^; to address donor-level confounding, we additionally computed per-donor combined-kernel transition mass and contrasted Memory B cell to ABC against Naïve B cell to ABC across n = 52 donors with age data: Switched-Memory to ABC averaged 0.059 versus Naïve to ABC 0.009 (**Supplementary Table 4**). Direction and statistical significance were preserved under every q-score weighting variant, demonstrating that the ageing Memory to ABC observation is not a tautological consequence of the q-score (**Fig. 6A, Supplementary Table 5**). Transition rates were near zero in donors under 20 years, rose gradually through middle age, and reached the highest values in donors over 60 years, with substantial inter-individual variation within each age group (**Fig. 6A)**. The progressive increase in unswitched Memory B to DN2 like B cell transition rates with age suggests that IgM^+^ unswitched memory B cells become increasingly prone to extrafollicular differentiation into DN2 like B cell state as individuals age, consistent with the increased frequency of DN2/AtM and ABC cells observed in aged cohorts^9,11^. To confirm that these per-donor transition probability differences reflect genuine biological variation rather than technical confounders, we compared total UMI count, number of detected genes, and mitochondrial read fraction between all pairwise population comparisons. Cohen’s *d* was ≤0.2 for all three covariates across all comparisons, and all 95% confidence intervals spanned zero, confirming that the transition probability analyses are not confounded by systematic differences in sequencing depth or cell quality between source populations(**Supplementary Fig. 5A**). Memory B cells transitioning to DN2 showed significantly elevated EF score, *ZEB2* expression, and fewer SHMs relative to non-transitioning memory cells with age (p < 0.05 each; **Supplementary Fig. 5B**).

**Figure 6.**
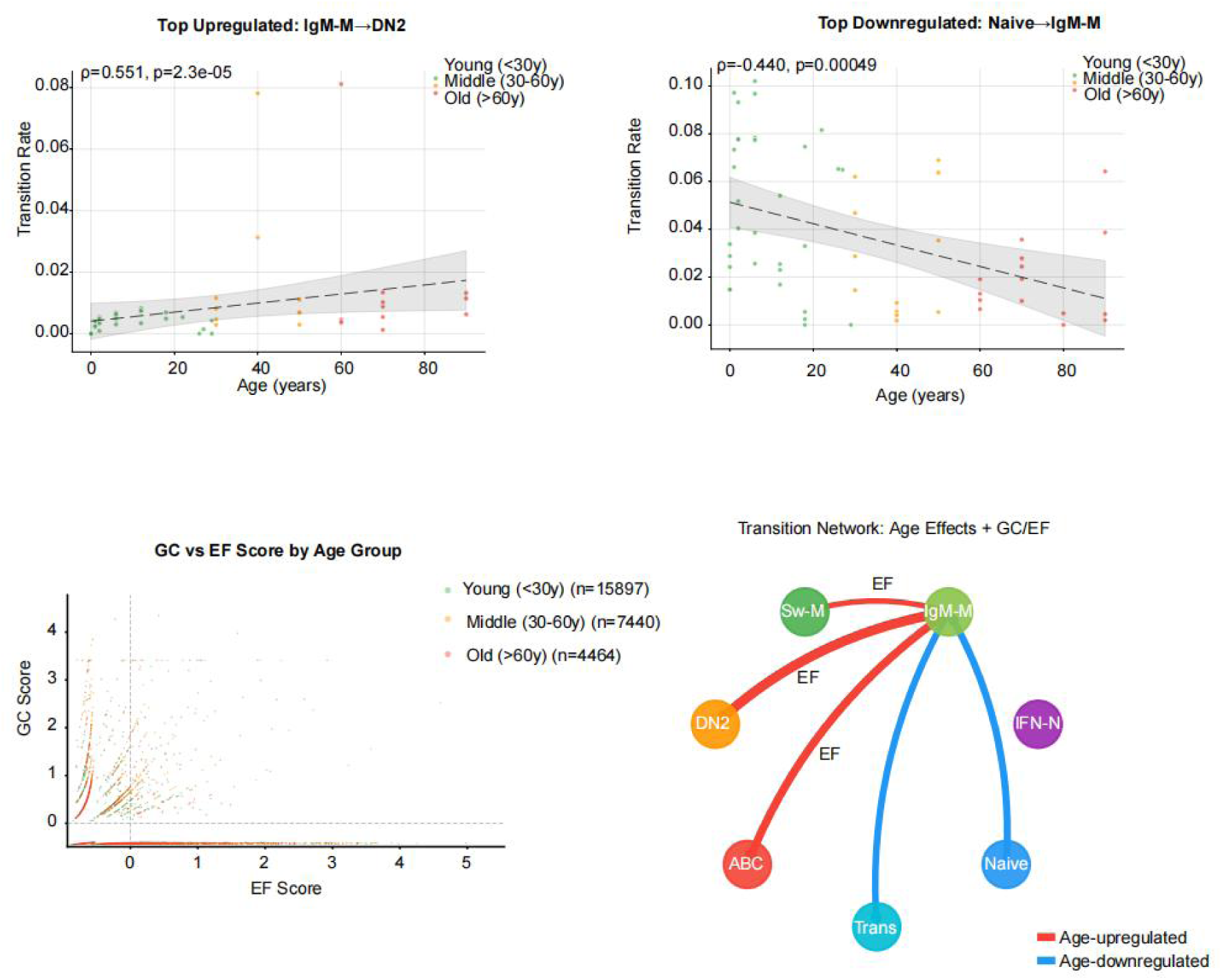
Age-dependent B cell transition upregulation is driven by the extrafollicular pathway. (A) Per-donor scatter plot of the IgM Memory to DN2 transition (Spearman ρ = 0.551, p = 2.3×10⁻⁵; fold change = 3.76, BH-corrected p = 0.028). Each dot represents one donor (n = 52), coloured by age group (green: young <30 y; orange: middle 30–60 y; red: old >60 y). Dashed line shows linear regression with 95% confidence interval. (B) Per-donor scatter plot of the Naïve Resting to IgM Memory transition (Spearman ρ = −0.440, p = 4.9×10⁻⁴; fold change = 0.38). (C) EF score versus GC score per cell, coloured by cell type. Young: green, middle age: yellow, old: red. (D) Network schematic of statistically significant age-dependent transitions. Red edges: age-upregulated; blue edges: age-downregulated. Edge width is proportional to |log₂(fold change)|.

The transition from Naïve Resting B cells to IgM^+^ Memory was significantly age-downregulated (Spearman ρ = −0.440, p = 4.9×10^−4^; **Fig. 6B**). Healthy ageing ABCs and SLE DN2 cells shared a number of TFs, and EF pathway markers (**Supplementary Fig. 5C**). In addition, the *FCRL5* and *FCER2* are positively correlated with ABC fate in both SLE cohorts and ageing cohorts (>60 years old, **Supplementary Fig. 5D**). This demonstrates that *FCRL5* and *FCER2* as a cross-dataset, cross-disease marker of extrafollicular B cell differentiation. GC and EF score analysis confirmed that the IgM^+^ B cell maturation in the older populations is EF dominated with lower GC score (**Fig. 6C**). Integrating the age-upregulated and age-downregulated transitions with the EF/GC pathway classification produced a unified transition network of age effects on B cell fate allocation (**Fig. 6D**). IgM^+^ Memory emerged as the hub of the age-sensitive network, with three age-upregulated extrafollicular transitions emanating from it: IgM^+^Memory B to DN2 (EF, red), IgM^+^Memory B to ABC (EF, red), and IgM^+^Memory B to Switched Memory (red). The single most prominent age-downregulated transition was Naïve to IgM^+^Memory B (blue) (**Fig. 6D**).

This network architecture suggests that ageing progressively reorients the B cell trajectory landscape around the IgM^+^ Memory B cell compartment, with IgM^+^ Memory B cells in older individuals increasingly maturing into DN2 and ABC via extrafollicular differentiation.

## 4. Discussion

By incorporating BCR maturation as a directional refinement of transcriptional trajectory inference, ClonoTrace resolves developmental transitions that RNA-only methods cannot reliably order. The q-score is a composite metric encoding isotype switching, SHM burden, and clonal expansion, which provides an independent temporal axis. The q-score constrains the Markov transition kernel and reinforces forward transitions and penalises biologically implausible reversals. Applied to peripheral blood B cells, ClonoTrace identified memory B cell extrafollicular differentiation route as a predominant route generating pathogenic DN2 and ABCs, and revealed the DN2 as the transition state for age-dependent rewiring of B cell maturation.

ClonoTrace identified that memory B cell extrafollicular re-differentiation, in addition to de novo activation from naïve precursors, constitutes the alternative pathway generating DN2 and ITGAX⁺ AtM cells in peripheral blood of SLE patients. Up-regulation of *ZEB2* and *RHOB* in bridge cells, accompanied by a decline in *BACH2* along the Memory to DN2 pseudotime trajectory, marks this axis, with *ZEB2* directly inducing *ITGAX* and *FCRL5* expression while suppressing CXCR5-mediated GC entry^8,20^. The lower SHM burden in DN2-committed cells suggests a shorter or none GC maturation, and BCR clonotype sharing between switched memory and DN2 populations. These findings help resolve the two competing hypothesis^11,12^. Both memory B cell and naïve B cell routes are present, but the memory pathway may be more activated in the peripheral blood of some SLE patients.

Three three pathways to age-associated B cells (ABC) were identified by ClonoTrace that are composed of naïve-to ABC, switched memory B cells to ABC, and IgM^+^ memory to ABC. Age-dependent rewiring of these transition pathways indicates that the B cell differentiation landscape undergoes systematic reconfiguration with advancing age. The IgM^+^ memory B cell to ABC transition emerges as the most age-sensitive pathway, with transition probability increasing substantially in older donors. This selective amplification suggests that ageing preferentially activates the IgM^+^ memory compartment for extrafollicular re-differentiation with lower antibody-secreting ability, rather than uniformly expanding all DN2 precursor populations. These age-dependent changes offer a candidate mechanistic framework for the well-documented expansion of T-bet⁺CD11c⁺ B cells in aged mice and elderly humans^9,10,10,14,20,21^.

ClonoTrace provides a quantitative framework for dissecting how therapeutic interventions reshape B cell transition landscapes at single-cell resolution. Monitoring DN2 percentage before and after anti-BAFF or anti-IFNAR1 therapy could test whether the pathogenic mechanisms identified by ClonoTrace represent a therapeutic intervention. We note that the present data do not themselves establish therapeutic predictive value; this is offered as a testable hypothesis for future intervention studies rather than a demonstrated application. More broadly, the principle of using receptor maturation as a directional prior is not limited to B cells; analogous signals such as T cell receptor diversification could extend trajectory inference to other lineages where transcriptomic ordering alone is insufficient. By demonstrating that immunoglobulin evolution and transcriptomic state can be jointly modelled within a single probabilistic framework, ClonoTrace establishes a paradigm for immune receptor-informed developmental biology that bridges clonal identity with cellular fate.

### Limitations

Several limitations warrant consideration. The SLE application relies on a single cohort (n = 9 patients), and clinical metadata (SLEDAI, treatment status, demographics) were not available for this cohort, so interferon stratification was inferred from an expression-derived ISG-module score rather than clinical labels, and validation in independent lupus datasets will be necessary to confirm the generalizability of the inferred transition hierarchy. ClonoTrace requires paired scRNA-seq and scV(D)J sequencing data, limiting applicability to datasets generated with compatible capture protocols. The q-score isotype ordering assumes a canonical maturation hierarchy that may not hold in mucosal tissues where IgA class switching can precede GC entry. Finally, the inferred trajectories and convergence architecture remain computational predictions; functional validation through lineage tracing or perturbation experiments will be essential to confirm the directionality and obligate nature of the DN2 intermediate. Several caveats further bound the directional interpretation. First, all trajectory directionality reported here is inferred rather than directly observed: neither RNA velocity nor experimental lineage tracing was available for these cohorts, so our inference indicates, rather than proves, the Memory to DN2/ABC ordering. Second, external replication in GSE254176 is directional and small-n (six donors), and the largest per-donor ratios are inflated by near-zero Naïve-source transition masses; we therefore emphasise the denominator-stabilised median (13.4×; epsilon-padded 11.8×) and the unanimous 6/6 sign concordance rather than the raw ratio magnitudes. Finally, the BCR-aware kernel relies on AntiBERTy sequence embeddings, a learned representation whose behaviour on rare or atypical receptor sequences is not separately characterised here.

## Author Contributions

H.L and M.Z. contributed equally to this work. H.L, J.Z and X.C conceptualized and supervised the project. M.Z. and H.L developed the ClonoTrace computational framework and performed all trajectory analyses. M.Z. and H.L performed scRNA-seq data processing, cell-type annotation, and transcription factor analyses. H.L and M.Z. performed BCR repertoire preprocessing, q-score pipeline development, and benchmarking analyses. H.L and M.Z. contributed to data processing and figure preparation. B.Z. and Q.L. collected SLE patient samples and provided clinical expertise. H.L, J.Z and X.C. interpreted the data and wrote the manuscript with input from all authors.

## Data Availability

Publicly available datasets used in this study are accessible from their original publications: fetal B cell data (Suo et al., Science 2022; Jardine et al., Nature 2021), adult lymph node data (Kim et al., Nature 2022). The SLE PBMC scRNA-seq and scV(D)J-seq data generated in this study have been deposited in the NCBI Gene Expression Omnibus (GEO; accession number to be assigned upon acceptance).

## Code availability

Code and a reproducible example notebook are available to editors and reviewers on request and will be public at https://github.com/alice-svg/ClonoTrace on publication.

## Acknowledgements

We thank all patients who participated in this study and the clinical staff at Xiangya Hospital. This work was supported by National Key R&D Program of China (2023YFA1801400), National Natural Science Foundation of China (82388201).

---

**Supplementary Figure 1.**
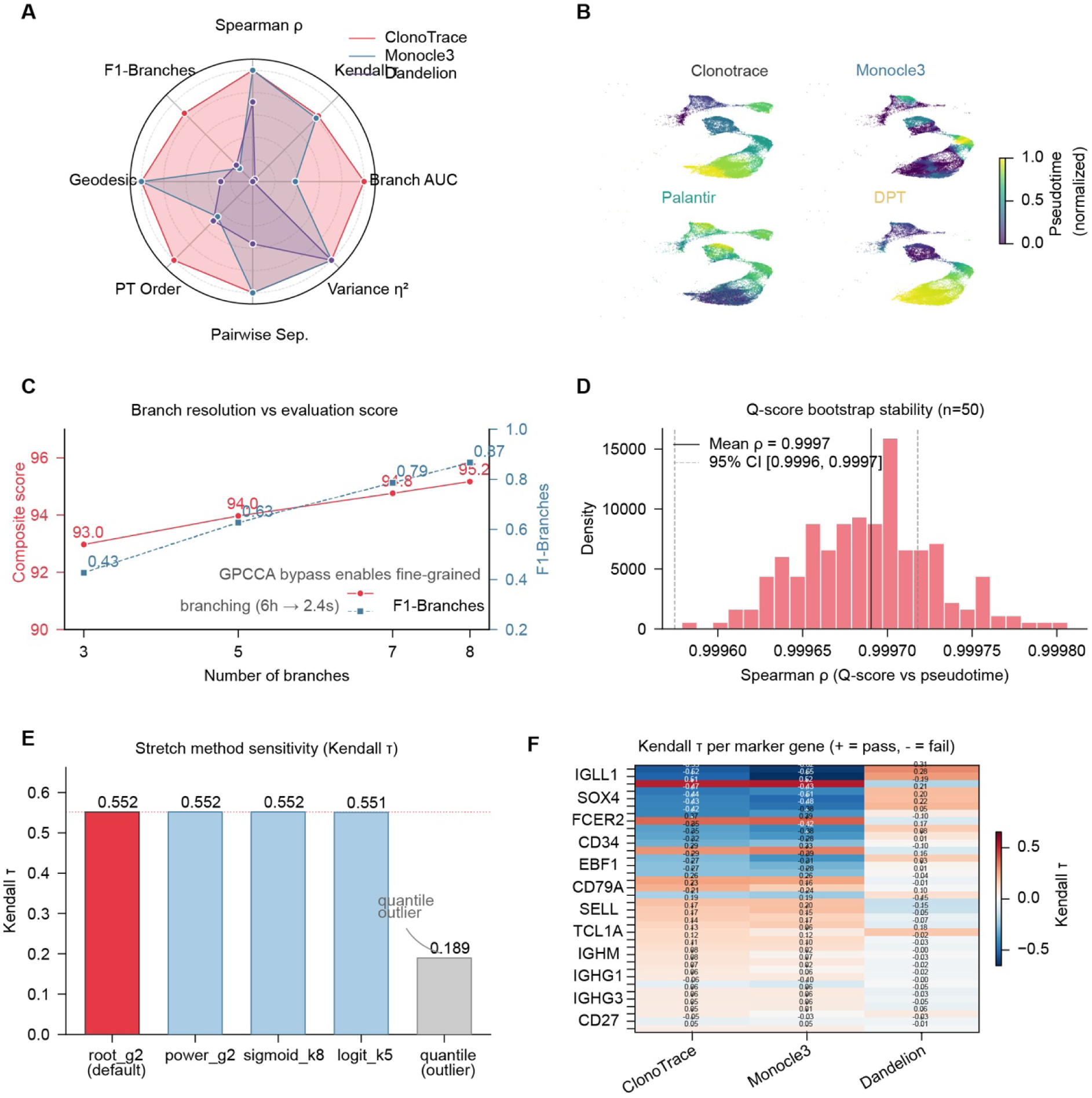
**(A)** Radar chart comparing ClonoTrace (red), Monocle 3 (light blue), and Dandelion (dark blue) across eight trajectory evaluation metrics: Spearman ρ, Kendall τ, Branch AUC, Variance η², PT Order, Pairwise Separation, Geodesic consistency, and F1-Branches. Each axis is scaled to the range [0,1] for visualization. **(B)** Side-by-side UMAP comparison of pseudotime gradients for ClonoTrace, Monocle 3, Palantir, and DPT. Color scale (0–1, normalized) is consistent across panels. ClonoTrace and Palantir produce globally ordered gradients; Monocle 3 shows partial ordering failures at intermediate stages; DPT compresses pseudotime in late-stage populations. **(C)** Branch resolution versus evaluation performance. Left y-axis (red): Composite score as a function of the number of branches resolved (n = 3–8). Right y-axis (blue dashed): F1-Branches score across the same range. Composite score increases from 93.0 (n = 3) to 95.2 (n = 8); **(D)** Bootstrap stability of q-score (n = 50 resampling replicates). Histogram of Spearman ρ between q-score and pseudotime across replicates. Solid vertical line indicates the mean (ρ = 0.9997); dashed lines indicate the 95% confidence interval [0.9996, 0.9997]. This panel and Supplementary Fig. 2D report the same stability metric — the Spearman ρ between the q-score computed on the full dataset and on independent 80% cell subsamples — on two different datasets (fetal here, GC in Supplementary Fig. 2D); the higher value here reflects the smaller, more homogeneous fetal dataset. **(E)** Sensitivity analysis of ClonoTrace trajectory performance to the choice of q-score contrast-stretching method. Kendall’s τ is shown for root_g2 (default, red, τ = 0.552), power_g2 (τ = 0.552), sigmoid_k8 (τ = 0.552), logit_k5 (τ = 0.551), and quantile (τ = 0.189, gray, labeled as outlier). **(F)** Kendall’s τ heatmap per marker gene for ClonoTrace, Monocle 3, and Dandelion across 13 representative canonical markers (*IGLL1, SOX4, FCER2, CD34, EBF1, CD79A, SELL, TCL1A, IGHM, IGHG1, IGHG3, CD27*). Positive τ (warm colors) indicates expected upregulation along pseudotime; negative τ (cool colors) indicates expected downregulation.

**Supplementary Fig 2.**
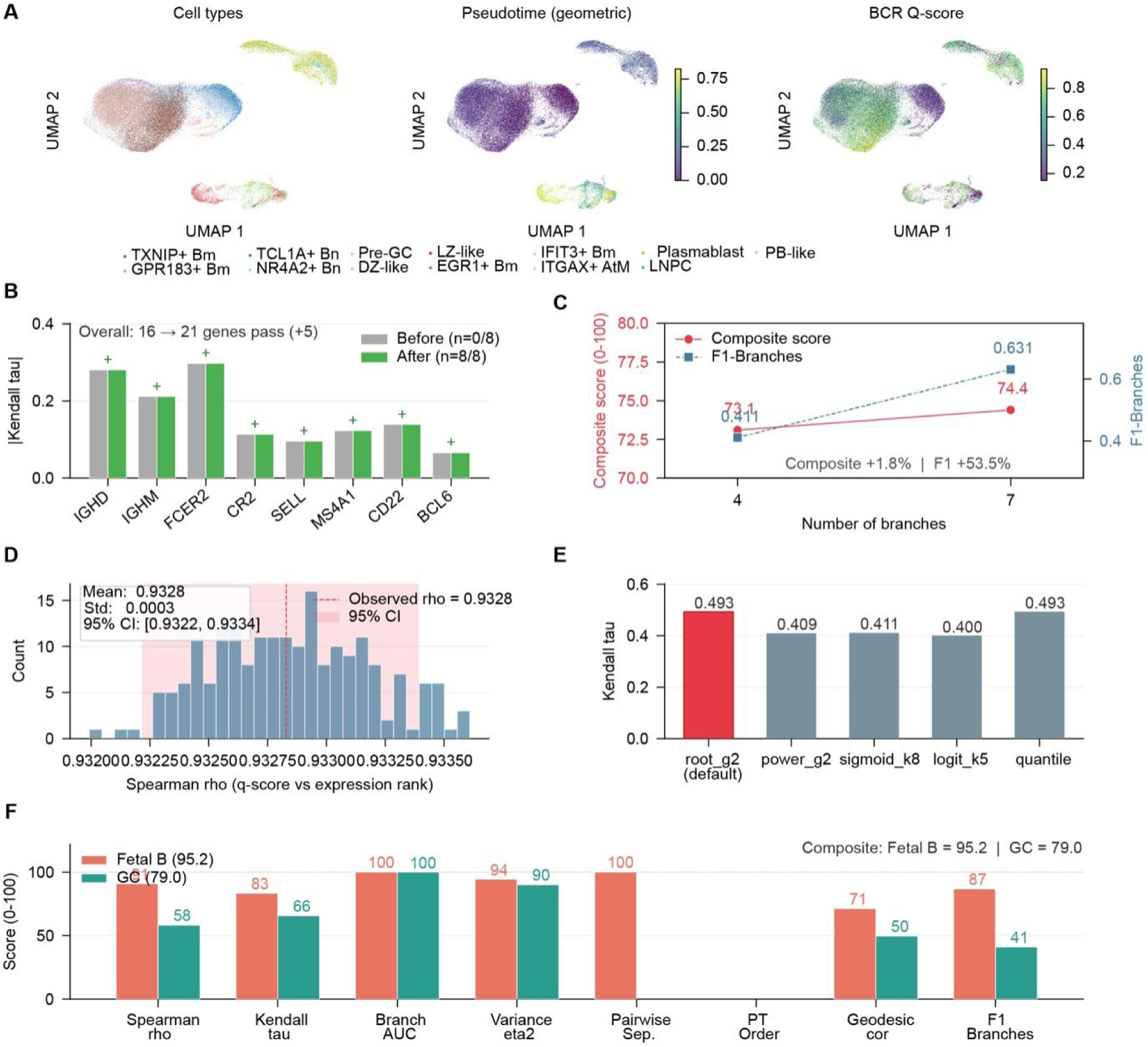
**(A)** UMAP embeddings of adult LN B cells colored by (left) cell type annotation, (center) ClonoTrace geometric pseudotime (0–1, normalized), and (right) BCR q-score (0–1). **(B)** Effect of BCR-aware pseudotime on naïve B cell marker gene directionality. **(C)** Branch resolution versus trajectory quality in the LN GC dataset. Left y-axis (red): Composite score across a branch-resolution sweep as branches increase from 4 (73.1) to 7 (74.4), a +1.8% improvement (the headline GC composite of 79.0 reported in the main text uses the corrected pipeline). Right y-axis (blue dashed) **(D)** Bootstrap stability of q-score in the LN GC dataset (n = 50 resampling replicates). Histogram of Spearman ρ between q-score and expression rank across replicates. Mean ρ = 0.9328, Std = 0.0003, 95% CI [0.9322, 0.9334]. This panel and Supplementary Fig. 1D report the same stability metric — the Spearman ρ between the q-score computed on the full dataset and on independent 80% cell subsamples — on two different datasets (GC here, fetal in Supplementary Fig. 1D); the lower value here reflects the larger, more heterogeneous GC atlas. **(E)** Sensitivity of LN GC trajectory performance to q-score contrast-stretching method. Kendall τ is shown for root_g2 (default, red, τ = 0.493), power_g2 (τ = 0.409), sigmoid_k8 (τ = 0.411), logit_k5 (τ = 0.400), and quantile (τ = 0.493). **(F)** Cross-dataset performance comparison between fetal B cells (Composite: 95.2; red) and LN GC B cells (Composite: 79.0; teal) across all eight evaluation metrics.

**Supplementary Figure 3.**
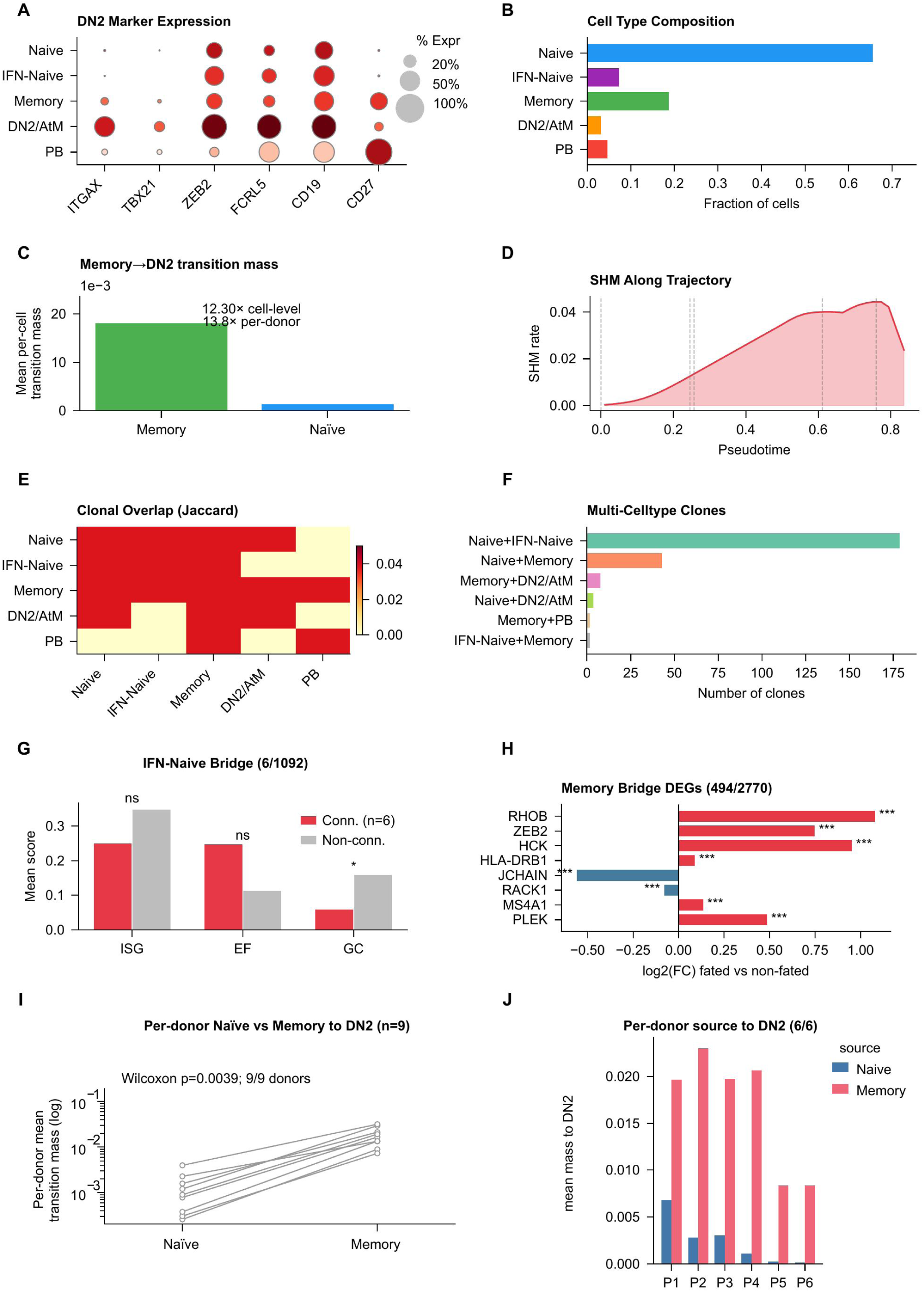
(**A**) Dot plot showing the expression of established DN2 B cell marker genes across five main peripheral blood B cell subsets identified from 9 SLE patients by paired scRNA-seq and scV(D)J-seq. (**B**) Bar plot showing the relative proportions of five main B cell subsets among total peripheral blood B cells. (**C**) Memory to DN2/AtM single-step transition mass; Memory exceeds Naïve 12.30-fold at the cell level and 13.8-fold as the per-donor mean, computed by ClonoTrace. (**D**) Line plot showing the mean SHM rate as a function of pseudotime. (**E**) Heatmap displaying the Jaccard index of clonal overlap between pairs of B cell subsets. The Jaccard index quantifies the proportion of shared clonotypes relative to the union of clonotypes in each pairwise comparison, with values ranging from 0 (no overlap, yellow) to 0.05 (maximal observed overlap, dark red). (**F**) Bar plot showing the number of expanded clones detected across two or more B cell subsets. (**G**) Bar plot comparing the mean module scores for interferon-stimulated genes (ISG), extrafollicular (EF), and germinal centre (GC) signature genes between DN2/AtM-fated bridge cells (Conn., n=6) and non-bridge IFN-Naïve cells (Non-conn., n=1,086). (**H**) Horizontal bar plot showing the log_2_fold change (log_2_FC) of differentially expressed genes between DN2/AtM-fated Memory bridge cells and non-fated Memory cells, identified by ClonoTrace bridge cells. (**I**) Per-donor paired Naïve versus Memory to DN2/AtM transition mass for all nine donors; every donor shows Memory above Naïve (paired Wilcoxon p = 0.0039). (**J**) Per-donor Memory and Naïve to DN2/AtM transition mass by patient.

**Supplementary Figure 4.**
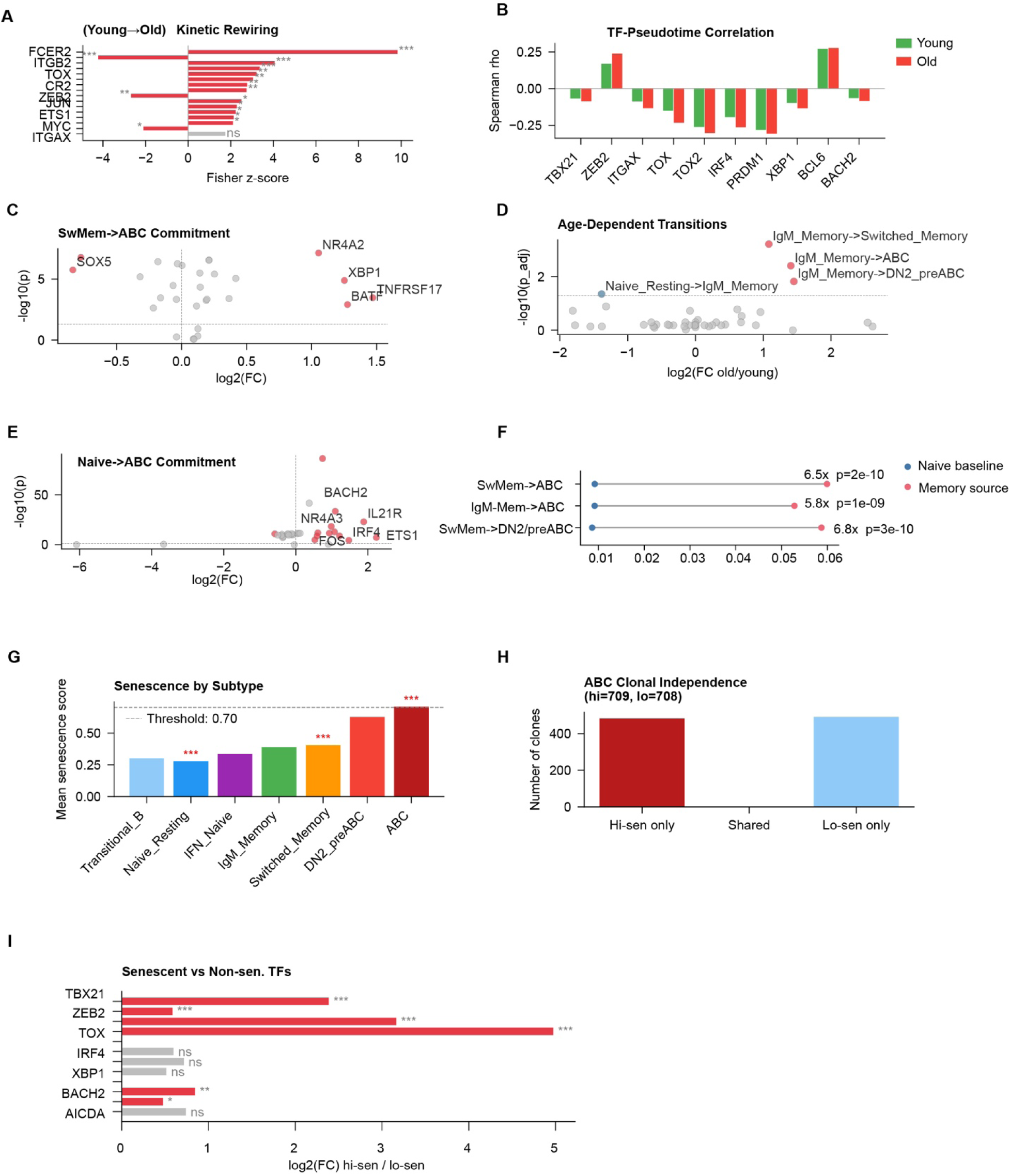
**(A)** Kinetic rewiring from young to old: Fisher z-scores quantifying the change in TF–pseudotime Spearman correlation between young (<30y) and old (>60y) donors for eight TFs **(B)** TF–pseudotime Spearman ρ for ten canonical TFs (*TBX21, ZEB2, ITGAX, TOX, TOX2, IRF4, PRDM1, XBP1, BCL6, BACH2*) in young (green bars) and old (red bars) donors. **(C)** Volcano plot of TF associations with SwMem to ABC commitment. **(D)** Volcano plot of age-dependent transition probability changes across all source–fate pairs. x-axis: log_2_FC (old/young); y-axis: −log_10_(adjusted p-value). **(E)** Volcano plot of TF associations with Naïve to ABC commitment (Naïve cells with high vs. low ABC fate probability). **(F)** Per-donor memory-source enrichment across 52 donors: Switched-Memory to ABC 6.5-fold, IgM-Memory to ABC 5.8-fold, and Switched-Memory to DN2_preABC 6.8-fold, all significant by paired Wilcoxon testing. **(G)** Mean senescence score by B cell subtype: Transitional B, Naïve Resting, IFN_Naïve, IgM_Memory, Switched_Memory, DN2_preABC, and ABC. The dashed line indicates the senescence threshold (score = 0.70). ABC cells significantly exceed the threshold (***; mean score ≈ 0.65–0.70). **(H)** ABC clonal independence by senescence group. Bar chart showing the number of clones containing only Hi-sen ABCs (709; dark red), shared clones containing both Hi-sen and Lo-sen ABCs (approximately 5; middle), and clones containing only Lo-sen ABCs (708; light blue). **(I)** TF expression comparison between senescent (Hi-sen) and non-senescent (Lo-sen) ABC cells. Bars show log_2_FC (Hi-sen / Lo-sen) for TBX21 (***), ZEB2 (***), ITGAX (***), TOX2 (***), IRF4 (ns), PRDM1 (ns), BCL6 (ns), MYC (ns), and AICDA (ns).

**Supplementary Figure 5.**
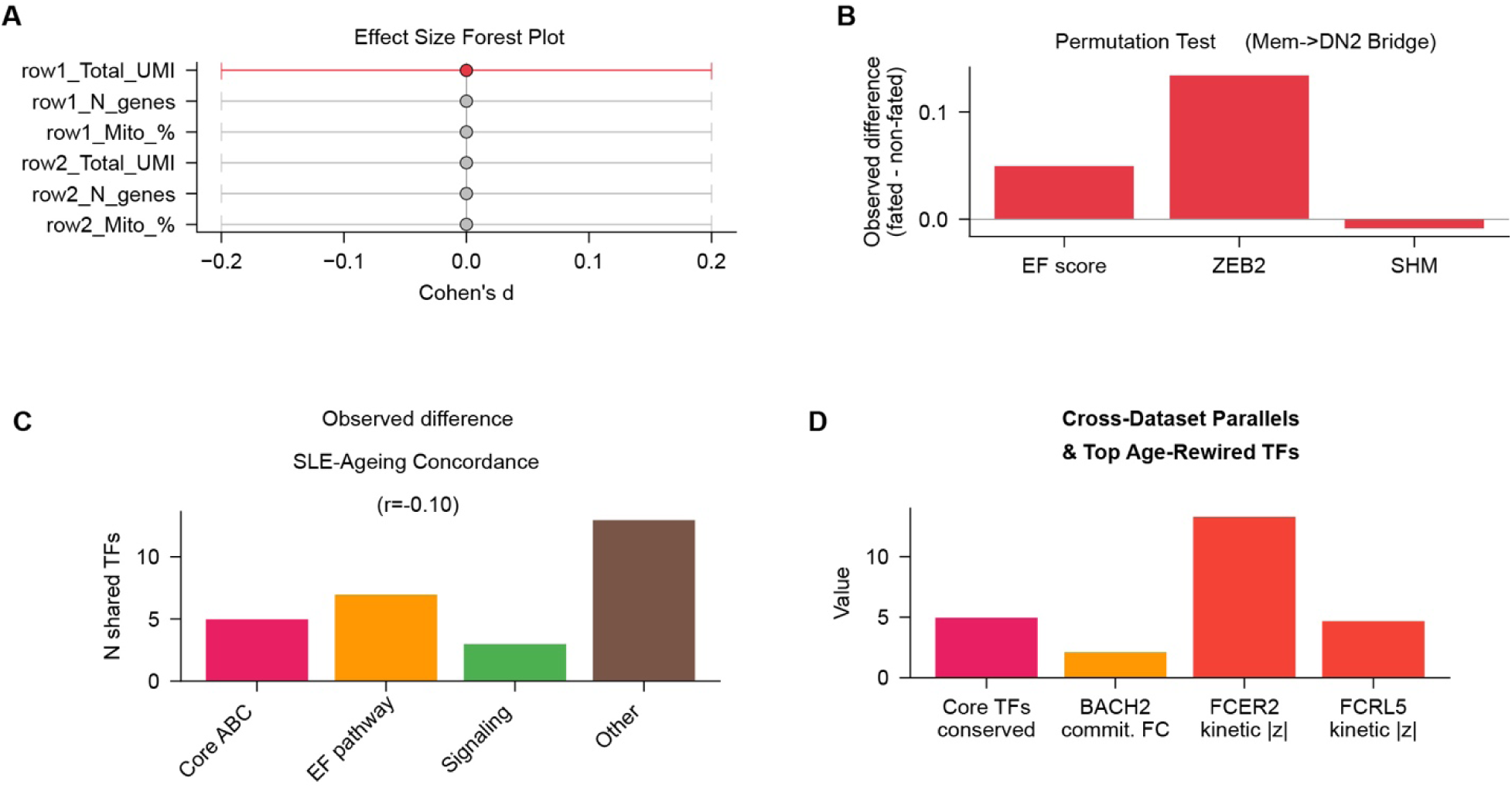
**(A)** Technical covariate balance. Effect-size forest plot (Cohen’s d) for row1/row2 Total UMI, gene counts, and mitochondrial percentages shows negligible effect sizes with 95% confidence intervals crossing zero, excluding technical confounding between groups. **(B)** Permutation test validating the extrafollicular (EF) origin of the memory-to-DN2 bridge. For each of three BCR maturation and EF identity features, EF score, *ZEB2* expression, and somatic hypermutation (SHM) burden. The observed group-mean difference between bridge-population cells (Mem→DN2 transitioning cells) and all other memory B cells is shown as a red bar, plotted against the null distribution derived from 1,000 random permutations of cell labels. **(C)** SLE–ageing TF concordance. The number of shared TFs significantly linked to ABC fate commitment (FDR < 0.05) is shown by functional category; the “Other” category contains the largest overlap, with an overall concordance of r = –0.10. **(D)** Cross-dataset validation of the BACH2, FCER2 and FCRL5 gatekeeper signal. Three quantitative metrics are shown: the number of core TFs conserved between the SLE and healthy-ageing ABC commitment signatures (Core TFs conserved); the Spearman ρ between BACH2 expression and ABC fate probability in the SLE PBMC cohort (BACH2 SLE ρ); and the log_2_ fold-change in FCER2 and FCRL5 pseudotime kinetics between young and old donors in the healthy-ageing cohort.

